# Olfactory responses of *Drosophila* are encoded in the organization of projection neurons

**DOI:** 10.1101/2022.02.23.481655

**Authors:** Kiri Choi, Won Kyu Kim, Changbong Hyeon

## Abstract

The projection neurons (PNs), reconstructed from electron microscope (EM) images of the *Drosophila* olfactory system, offer a detailed view of neuronal anatomy, providing glimpses into information flow in the brain. About 150 uPNs constituting 58 glomeruli in the antennal lobe (AL) are bundled together in the axonal extension, routing the olfactory signal received at AL to mushroom body (MB) calyx and lateral horn (LH). Here we quantify the neuronal organization by inter-PN distances and examine its relationship with the odor types sensed by *Drosophila*. The homotypic uPNs that constitute glomeruli are tightly bundled and stereotyped in position throughout the neuropils, even though the glomerular PN organization in AL is no longer sustained in the higher brain center. Instead, odor-type dependent clusters consisting of multiple homotypes innervate the MB calyx and LH. Pheromone-encoding and hygro/thermo-sensing homotypes are spatially segregated in MB calyx, whereas two distinct clusters of food-related homotypes are found in LH in addition to the segregation of pheromone-encoding and hygro/thermo-sensing homotypes. We find that there are statistically significant associations between the spatial organization among a group of homotypic uPNs and certain stereotyped olfactory responses. Additionally, the signals from some of the tightly bundled homotypes converge to a specific group of lateral horn neurons (LHNs), which indicates that homotype (or odor type) specific integration of signals occurs at the synaptic interface between PNs and LHNs. Our findings suggest that before neural computation in the inner brain, some of the olfactory information are already encoded in the spatial organization of uPNs, illuminating that a certain degree of labeled-line strategy is at work in the *Drosophila* olfactory system.

## Introduction

Anatomical details of neurons obtained based on a full connectome of the *Drosophila* hemisphere reconstructed from EM image datasets (***Bates et al., 2020***; ***Scheffer et al., 2020***) offer the wiring diagram of the brain, shedding light on the origin of brain function. Out of the immense amount of data, we study the second-order neurons, known as the projection neurons (PNs) of the olfactory system. It is the PNs that bridge the olfactory receptor neurons (ORNs) in the antenna and maxillary palp to higher olfactory centers where neural computation occurs for *Drosophila* to sense and perceive the environment (***Hallem and Carlson, 2004***). The three neuropils, namely the antennal lobe (AL), mushroom body (MB) calyx, and lateral horn (LH), are the regions that abound with an ensemble of axonal branches of PNs and synapses (Figure 1). PNs can be classified as uniglomerular and multiglomerular PNs based on their structure and connectivity to other PNs. The uniglomerular PNs (uPNs) in AL constitute glomeruli that collect olfactory signals from ORNs of the same receptor type (***Gao et al., 2000***; ***Couto et al., 2005***). uPNs innervating MB calyx and LH relay the signals further inside the brain through synaptic junctions with the Kenyon cells (KCs) and lateral horn neurons (LHNs), respectively. Multiglomerular PNs (mPNs), on the other hand, innervate multiple glomeruli, regulating the signals from ORNs and often contributing to inhibitory regulation (***Berck et al., 2016***). PNs can functionally be categorized into either excitatory (cholinergic) or inhibitory (GABAergic), where a many GABAergic PNs tend to innervate only one of the two higher olfactory centers (***Schultzhaus et al., 2017***; ***Shimizu and Stopfer, 2017***).

**Figure 1.**
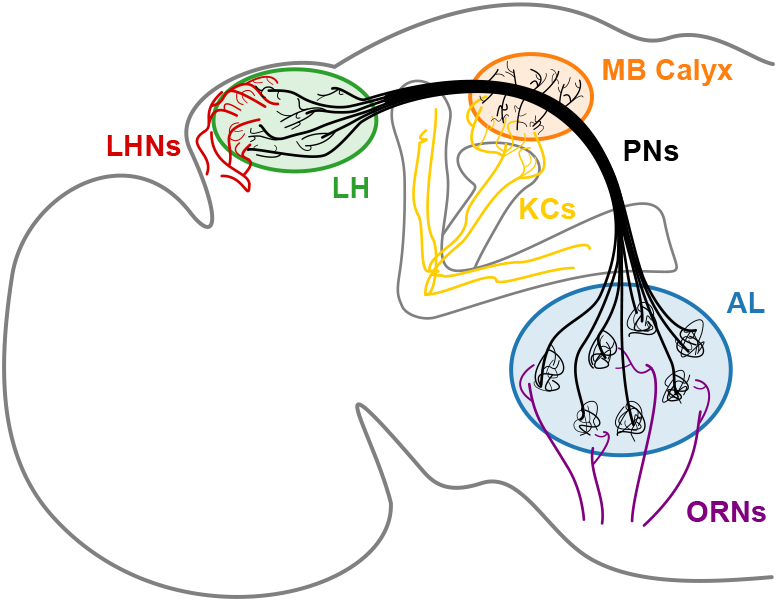
A schematic of the *Drosophila* olfactory system. uPNs comprising each glomerulus in AL collect input signals from ORNs of the same receptor type and relay the signals to MB calyx and LH. uPNs in MB calyx synapse onto KCs; and uPNs in LH synapse onto LHNs.

Since the seminal work by Ramón y Cajal (***y Cajal, 1911***), who recognized neurons as the basic functional units of the nervous system, there have been a series of attempts at classifying neurons using different representations of neuronal morphologies and at associating the classified anatomies with their electrophysiological responses and functions (***Uylings and Van Pelt, 2002***; ***Scorcioni et al., 2008***; ***Jefferis et al., 2007***; ***Seki et al., 2010***; ***Gillette and Ascoli, 2015***; ***Lu et al., 2015***; ***Li et al., 2017***; ***Kanari et al., 2018***; ***Mihaljević et al., 2018***; ***Gouwens et al., 2019***; ***Laturnus et al., 2020***). Systematic and principled analyses of neuronal anatomy would be a prerequisite for unveiling a notable link between the PN organization and olfactory representations. Several different metrics involving spatial projection patterns (***Jefferis et al., 2007***), electrophysiological properties (***Seki et al., 2010***; ***Gouwens et al., 2019***), topological characteristics (e.g. morphometrics) (***Uylings and Van Pelt, 2002***; ***Scorcioni et al., 2008***; ***Lu et al., 2015***; ***Mihaljević et al., 2018***; ***Gouwens et al., 2019***), intersection pro*file*s (***Gouwens et al., 2019***), and NBLAST scores (***Jeanne et al., 2018***; ***Zheng et al., 2018***; ***Bates et al., 2020***; ***Scheffer et al., 2020***) have been utilized in the past. More recently, machine learning approaches have been popularized as a tool for classification tasks (***Vasques et al., 2016***; ***Buccino et al., 2018***; ***Mihaljević et al., 2018***; ***Zhang et al., 2021***).

Among a multitude of information that can be extracted from the neural anatomy associated with uPNs, the inter-PN organization draws our attention. To compare spatial characteristics of uPNs across each neuropil and classify them based on the odor coding information, we confine ourselves to uPNs innervating all three neuropils, most of which are cholinergic and follow the medial antennal lobe tract (mALT). Within this scope, we first calculate inter-PN distance matrices in each neuropil and study them based on the glomerular types (homotypes) to discuss how the inter-PN organization changes as the PNs extend from AL to MB calyx and from AL to LH. We have conducted statistical analyses to unravel potential associations between the uPN organization and the behavioral responses of *Drosophila* to external stimuli encoded by glomerular homotypes, finding that certain odor types and behavioral responses are linked to a characteristic inter-neuronal organization. The map of synaptic connectivity between uPNs and the third-order neurons (KCs and LHNs in MB calyx and LH, respectively) complements the functional implication of the association between the inter-PN organization and olfactory processing. We discover that for some odor types which demand fast responses from the organism, the *Drosophila* olfactory system leverages the efficiency of the labeled-line design in sensory information processing (***Min et al., 2013***; ***Howard and Gottfried, 2014***; ***Andersson et al., 2015***; ***Galizia, 2014***).

## Results

### Spatial organization of neurons inside neuropils

First, we define a metric with which to quantify the spatial proximity between neurons. Specifically, the inter-PN distance *d*_*αβ*_ is the average taken over the minimum Euclidean distances between two uPNs *α* and *β*, such that *d*_*αβ*_ is small when two uPNs are tightly bundled together (see Equation 1 and Figure S1A). Although metrics such as the NBLAST score (***Costa et al., 2016***) and others (***Kohl et al., 2013***) can be used to study the PN organization, these metrics take both the morphological similarity and the spatial proximity into account. Therefore, the features of PN organization captured by the NBLAST distance are not necessarily aligned with *d*_*αβ*_ (see Figure S1B). In this study, we have deliberately chosen the metric *d*_*αβ*_ instead of the NBLAST score as we are only interested in the spatial proximity between two neurons.

The distances *d*_*αβ*_ (Equation 1) between all the possible pairs (*α* and *β*) of 135 uPNs are visualized in the form of a matrix (Figure 2). We perform hierarchical clustering on the distance matrix for uPNs in each neuropil (see the outcomes of *d*_*αβ*_-based clustering analysis in Figure S2 and Methods for the details). Individual clusters from the hierarchical clustering of uPNs in MB calyx and LH are visualized in Figures 3 and 4 with the colors denoting the odor types encoded by the individual uPNs, which will be discussed in detail later.

**Figure 2.**
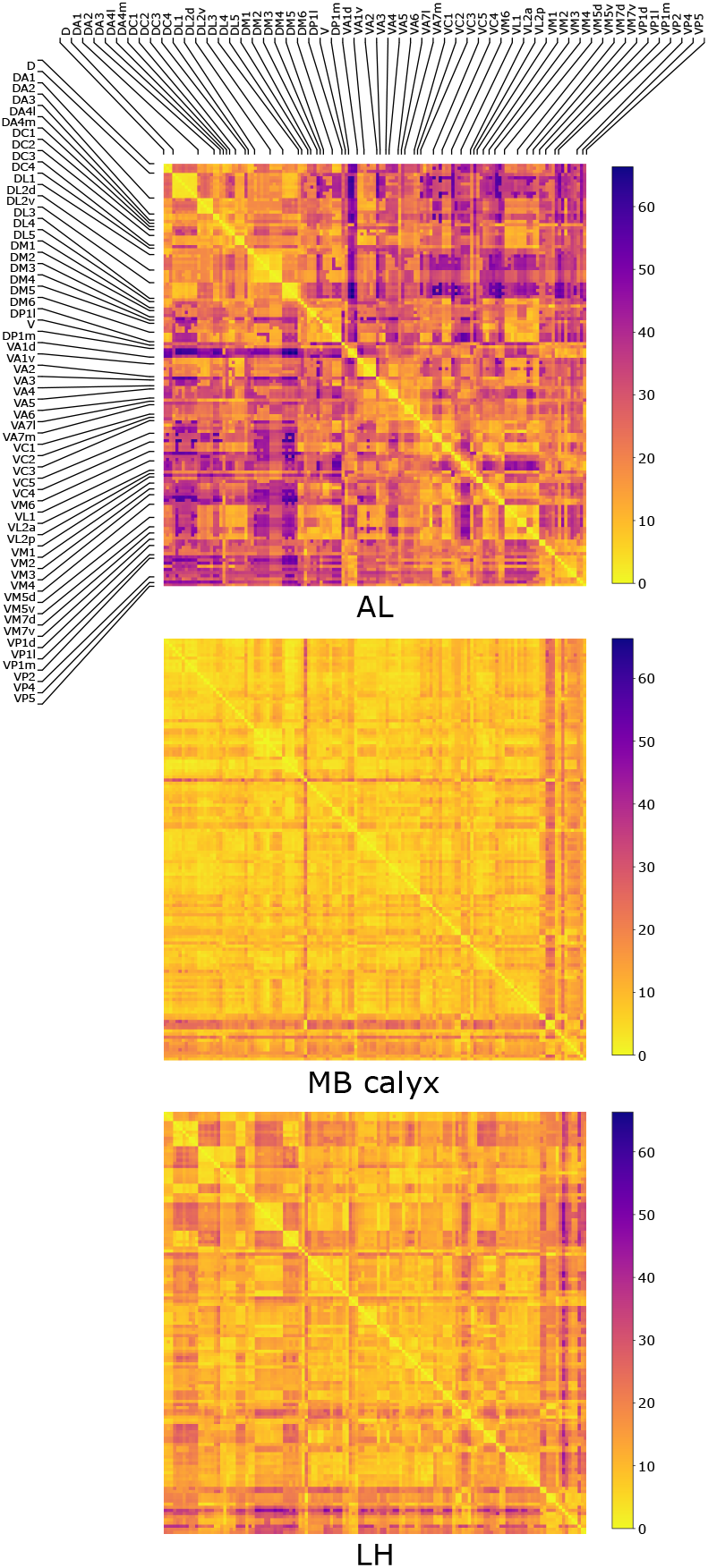
The three matrices representing the pairwise distances *d*_*αβ*_ in units of *μ*m between individual uPN in AL, MB calyx, and LH. The diagonal blocks represent the homotypic uPNs comprising the 57 glomerular homotypes available in the FAFB dataset (***Bates et al., 2020***), labeled at the edges.

**Figure 3.**
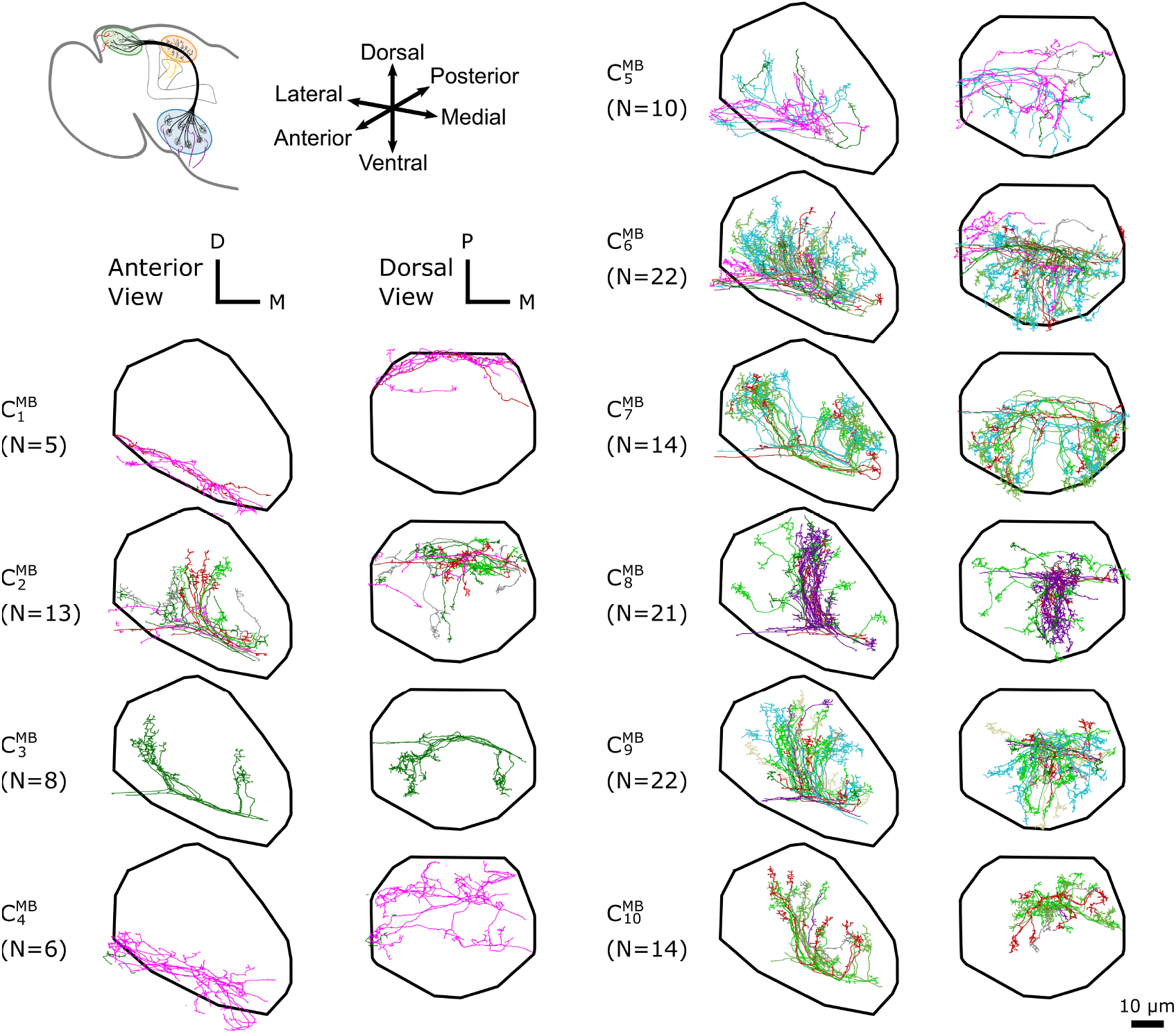
The *d*_*αβ*_ -based clustering on uPNs in MB calyx resulting in 10 clusters. The individual uPNs are color-coded based on the encoded odor types (Dark green: decaying fruit, lime: yeasty, green: fruity, gray: unknown/mixed, cyan: alcoholic fermentation, red: general bad/unclassified aversive, beige: plant matter, brown: animal matter, purple: pheromones, pink: hygro/thermo) (***Mansourian and Stensmyr, 2015***; ***Bates al., 2020***). The first and second columns illustrate the anterior and the dorsal view, respectively (D: dorsal, M: medial, P: posterior). The black line denotes the approximate boundary of MB calyx.

**Figure 4.**
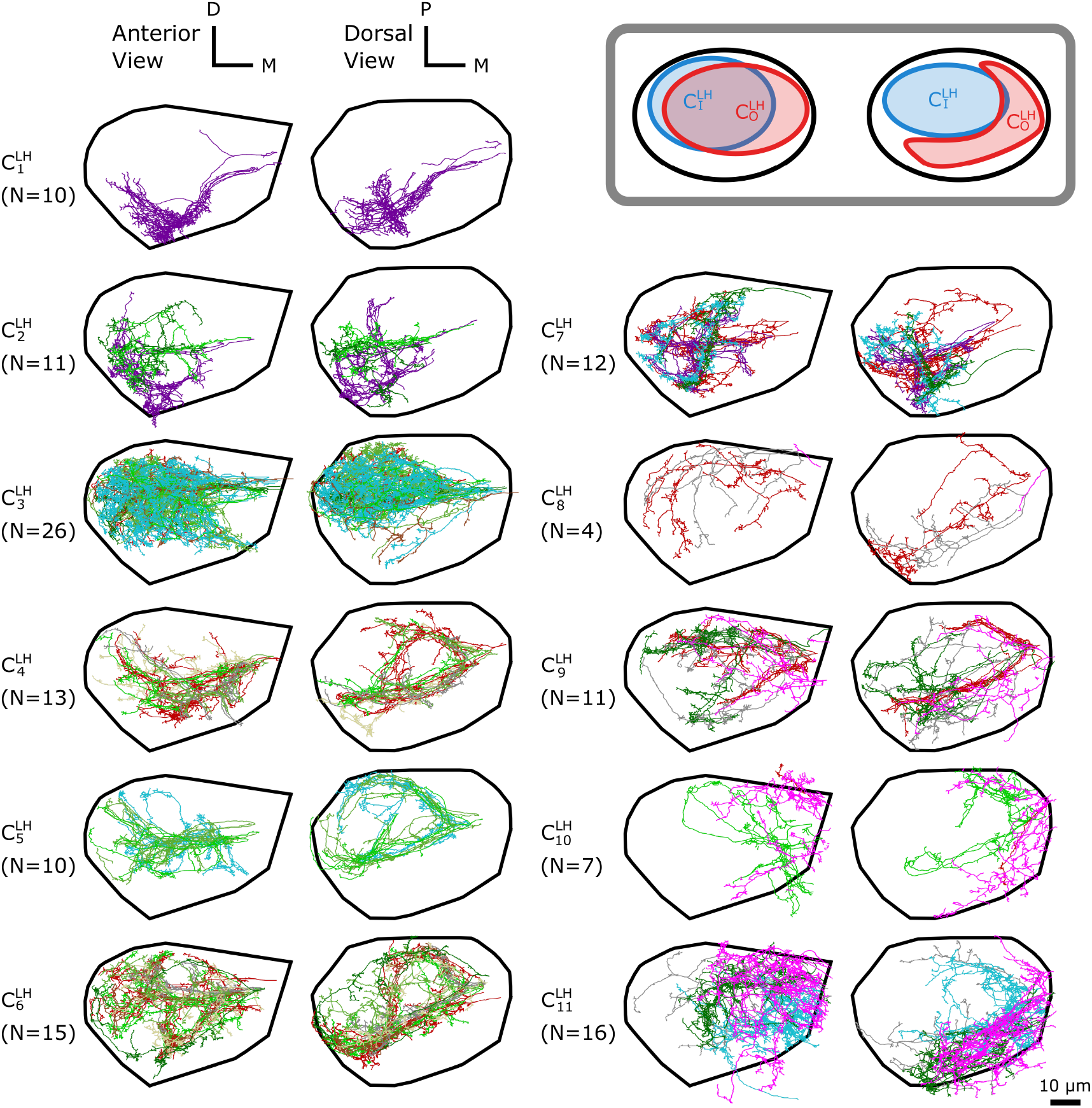
The *d*_*αβ*_-based clustering on uPNs in LH resulting in 11 clusters. (inset) A cartoon illustrating the relative position between clusters 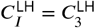 and 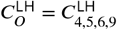. The individual uPNs are color-coded based on the encoded odor types (Dark green: decaying fruit, lime: yeasty, green: fruity, gray: unknown/mixed, cyan: alcoholic fermentation, red: general bad/unclassified aversive, beige: plant matter, brown: animal matter, purple: pheromones, pink: hygro/thermo). The first and second columns illustrate the anterior and the dorsal view, respectively (D: dorsal, M: medial, P: posterior). The black line denotes the approximate boundary of LH.

In MB calyx, the hierarchical clustering divides the uPNs into 10 clusters (Figure 3). Clusters 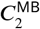 and 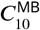 largely encompass the posterior region and clusters 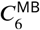 and 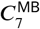 encompass the anterior region of the neuropil. The cluster 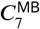 shows a characteristic biforked pattern projecting to the lateral and medial regions. The cluster 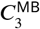 also exhibits the same structural pattern but is composed of a tight bundle of uPNs that are part of DL2d and DL2v. The cluster 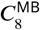 is located between the biforked innervation pattern of clusters 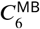 and 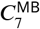, and predominantly innervates the dorsal region. Lastly, clusters 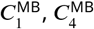, and 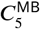, innervate the ventral region of MB calyx, spatially separated from other uPNs.

In LH, 11 clusters are identified (Figure 4). The cluster 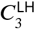 is the largest, which mainly innervates the dorsal posterior region of LH. Clusters 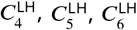, and 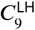 display variable biforked projection patterns along the horizontal plane, enveloping the boundary of the cluster 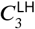. This creates a spatial pattern where a large blob of uPNs 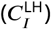 are surrounded by a claw-like structure 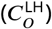 (Figure 4, inset). Clusters 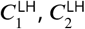, and 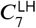 innervate the anterior-ventral region and display clear segregation from the other uPNs. Another group composed of clusters 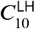 and 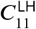 innervates the anterior-dorsal-medial region.

We use Pearson’s *x*^2^-test (see Methods for the details) to assess the likelihood of dependence between the *d*_*αβ*_-based clustering outputs for MB calyx, LH, and the glomerular labels (homotypes) statistically significant correlations are found in terms of both the p-value and the Cramér’s *V* (see Table S1 and Methods for a detailed explanation of the meaning behind the p-value and the Cramér’s *V*), the latter of which is analogous to the correlation coefficient for the *x*^2^-test. The mutual information between the same set of nominal variables, which is calculated to verify our *x*^2^-tests (see Methods), offers a similar conclusion (see Supplementary Information and Table S2).

We also categorize the spatial organization of uPNs in reference to the glomerular labels. The homotypic uPNs constituting a tightly bundled glomerulus in AL manifest themselves as the block diagonal squares in the *d*_*αβ*_-matrix (Figure 2). This is apparent in the dendrogram constructed from the distance matrix for the uPNs at AL (Figure S3A), where uPNs sharing the same glomerular label are grouped under a common branch, thereby demonstrating the spatial proximity between uPNs forming the same glomerulus. The *d*_*αβ*_-matrix indicates that such organizations are also preserved in MB calyx and LH. However, clear differences are found in the off-diagonal part of *d*_*αβ*_ matrices (Figure 2).

To conduct a quantitative and concise analysis of *d*_*αβ*_ matrices, we define the mean intra- and inter-homotypic uPN distances, 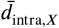 and 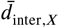 (see Methods for detailed formulation). The 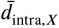 is the average distance between uPNs in the same homotype and measures the degree of uPNs in the homotype *X* being bundled. Therefore, a smaller 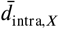 signifies a tightly bundled structure of *X*-th homotypic uPNs (see Figure S4 for raw 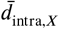 values). Similarly, 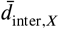, which measures the degree of packing (or segregation), is defined as the average distance between the neurons comprising the *X*-th homotype and neurons comprising other homotypes. Thus, a small value of 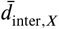 signifies tight packing of heterotypic uPNs around *X*-th homotype, while a large value indicates that the homotypic uPNs comprising the homotype *X* are well segregated from other homotypes (see Figure S4 for raw 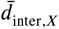 values).

The degrees of bundling averaged over all homotypes 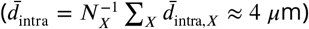 are comparable over all three neuropils (blue dots in Figure 5A). On the other hand, from 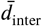, which is defined as the mean inter-homotype distance averaged over all *X*s, we find that homotypic uPNs are well segregated from others in AL as expected, whereas spatial segregation among homotypes is only weakly present in MB calyx (orange dots in Figure 5A and the cartoon of Figure 5B).

**Figure 5.**
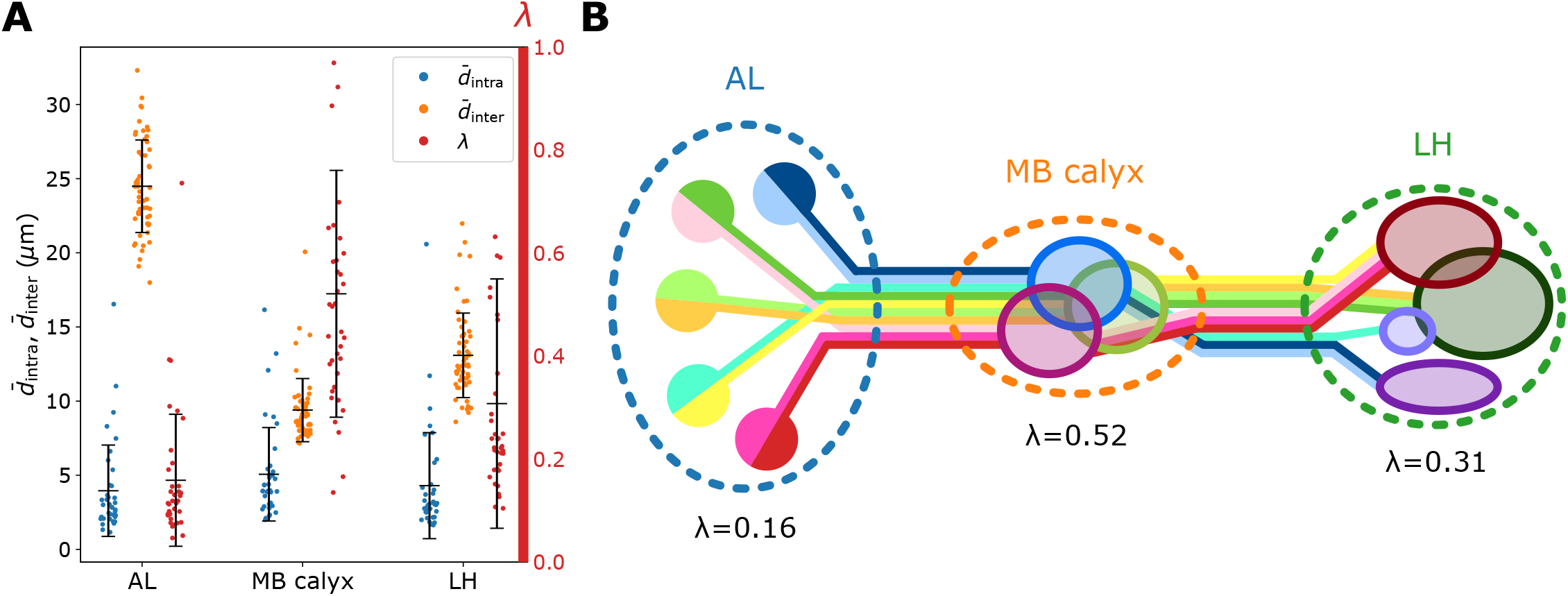
Organization of homotypic uPNs in the three neuropils. (A) A graph depicting 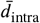 (blue, degree of bundling), 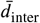 (orange, degree of packing), and the ratio between the two distances *λ* (red, degree of overlapping). Error bars depict the standard deviation. (B) Diagram illustrating the overall organization of uPNs at each neuropil. Homotypic uPNs are tightly bundled and segregated in AL. Several groups of homotypic uPNs form distinct heterotypic spatial clusters at higher olfactory centers, extensively overlapping in MB calyx (see Figure3).

Next, we take the ratio of mean intra- to inter-PN distances of *X*-th homotype as *λ*_*X*_ to quantify the degree of overlapping around *X*-th homotype (see Methods). The term *‘*overlapping*’* is specifically chosen to describe the situation where different homotypes are occupying the same space. A large value of *λ*_*X*_ (particularly *λ*_*X*_ > 0.4) suggests that the space occupied by the uPNs of the *X*-th homotype is shared with the uPNs belonging to other homotypes. The value 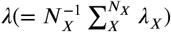 averaged over all the homotypes (red in Figure 5A) suggests that the extent of overlapping between uPNs is maximal in MB calyx and minimal in AL (Figure 5).

Figures 6A and 7 show individual values of *λ*_*X*_ for all homotypes in the three neuropils. We identify the following features: (i) In AL, *λ*_*X*_ ≤0.4 for all homotypes except DL5, indicating that homotypic uPNs are tightly bundled and segregated from uPNs in other glomeruli; (ii) In MB calyx, a large portion (≈65%) of *λ*_*X*_’s exceed 0.4 and even the cases with *λ*_*X*_ > 1 are found (VC5, DL5), implying that there is a substantial amount of overlap between different homotypes; (iii) Although not as significant as those in AL, many of uPNs projecting to LH are again bundled and segregated in comparison to those in MB calyx (see Figure 7B). (iv) The scatter plot of *λ*_*X*_ between MB calyx and LH (Figure 7C) indicates that there exists a moderate positive correlation (*r* = 0.642, *p* < 0.0001) between *λ*_*X*_ at MB calyx and LH (Figure 7C). This implies that a higher degree of overlapping in MB calyx carries over to the PN organization in LH.

**Figure 6.**
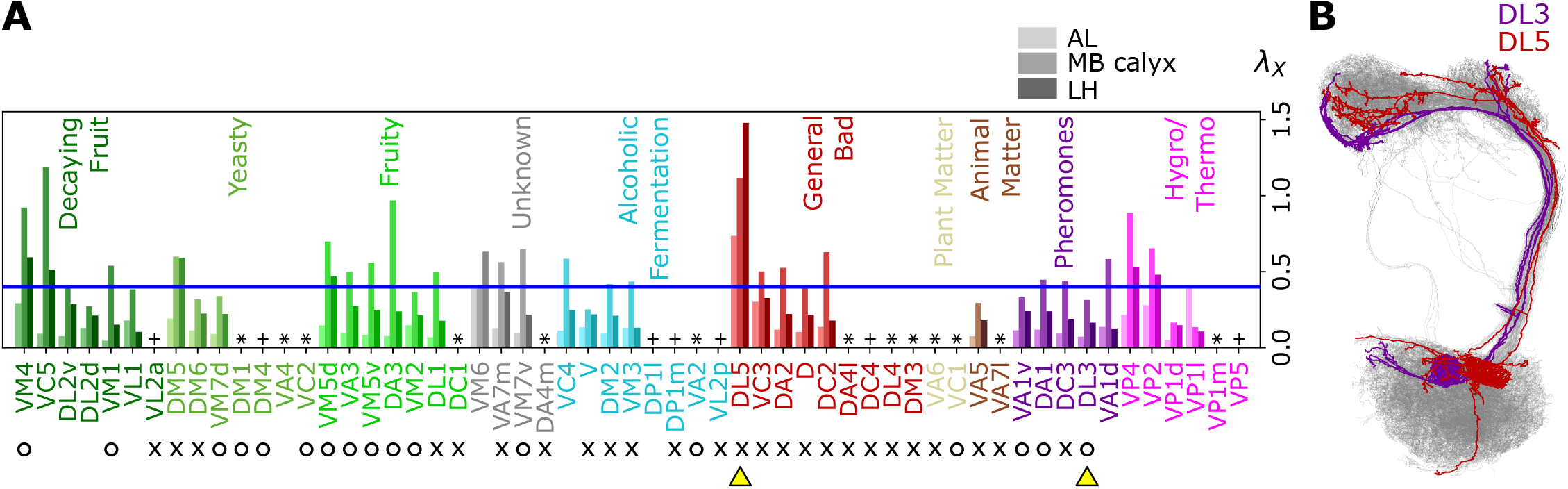
Degree of overlapping between inter-homotypic uPNs, *λ*_*X*_ (*X* = VM4, VC5, …, VP5). (A) The degree of overlapping (*λ*_*X*_) for *X*-th homotype in AL, MB calyx, and LH (from lighter to darker colors). The homotype label is color-coded based on the odor types associated with the glomerulus obtained from the literature and is sorted based on the value of *λ*_*X*_ for each odor type at LH. Asterisks (*) mark homotypes composed of a single uPN while plus (+) mark homotypes composed of a single uPN under our selection criterion but are actually a multi-uPN homotype, whose intra-homotype uPN distance is not available. O (attractive) and × (aversive) indicate the putative valence information collected from the literature. The blue horizontal line denotes *λ*_*X*_ = 0.4. (B) Two homotypes, DL3 (purple) and DL5 (red), which are indicated by yellow triangles in (A), are highlighted along with other uPNs (gray).

**Figure 7.**
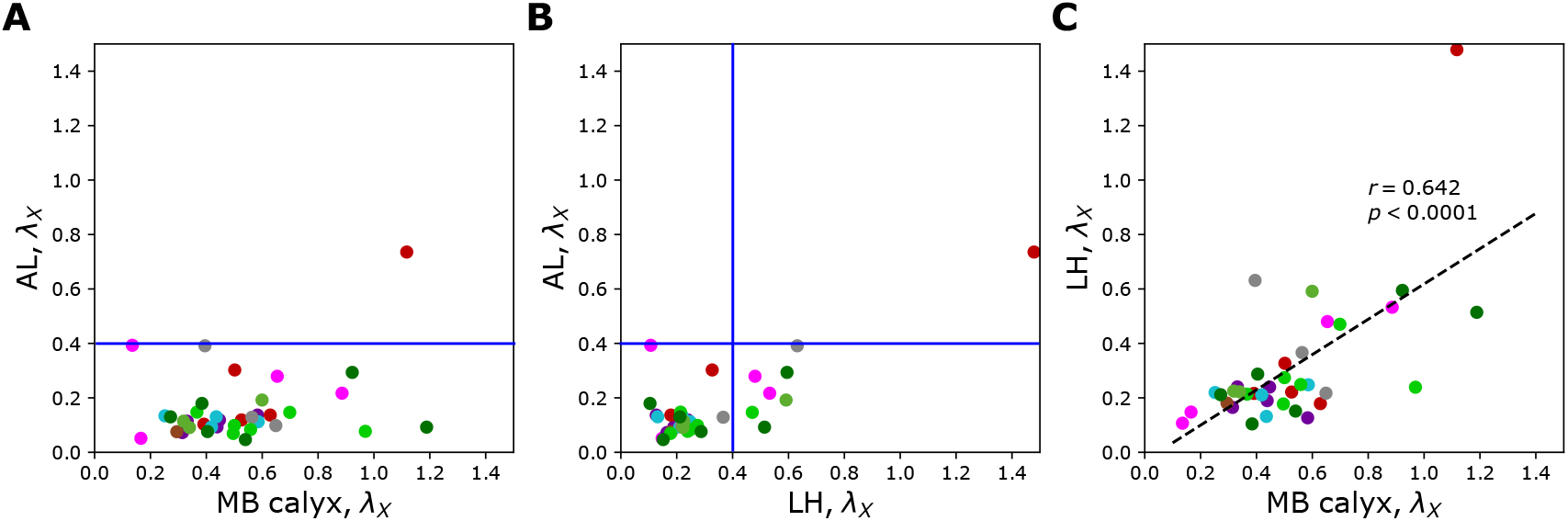
Scatter plots depicting the relationships between *λ*_*X*_ s at two different neuropils; (A) AL versus MB calyx, (B) AL versus LH, and (C) LH versus MB calyx. The color code is the same as in Figure 6A. The blue lines in (A) and (B) denote *λ*_*X*_ = 0.4.

The entire neuron morphologies of uPNs from two homotypes with a small (*X* = DL3) and a large (*X* = DL5) *λ*_*X*_ s in LH are visualized along with the other uPNs (gray) (Figure 6B). The homotype DL3, which seldom overlaps with others in AL (*λ*_DL3_ ≈ 0.07) and LH (*λ*_DL3_ ≈ 0.17), displays an increased overlapping in MB calyx (*λ*_DL3_ ≈ 0.31). Therefore, DL3 is tightly packed in AL and LH, whereas it is relatively dispersed in MB calyx. Meanwhile, the homotype DL5 displays a significant dispersion in all three neuropils, although the dispersion is the smallest in AL (*λ*_DL5_ ≈ 0.74) compared to that in MB calyx (*λ*_DL5_ ≈ 1.1) and LH (*λ*_DL5_ ≈ 1.5).

Taken together, the organization of olfactory uPNs varies greatly in the three neuropils. The clear homotype-to-homotype segregation in AL no longer holds in MB calyx. Instead, the *d*_*αβ*_-based clustering suggests the presence of clusters made of multiple different homotypic uPNs (Figure 5B). For some homotypes, the well-segregated organizations in AL are recovered when they reach LH (compare Figures 7A and B).

### Relationship between neuronal organization and olfactory features

Now we explore how the structural features identified from our clustering outputs are associated with odor types and valences (behavioral responses). As briefly mentioned earlier, the color codes in Figures 3, 4, 6, and 7 depict odor types encoded by corresponding homotypic uPNs, which follow the same categorical convention used by ***Mansourian and Stensmyr*** (***2015***) and ***Bates et al***. (***2020***). The O and × represent the putative valence, which indicates whether *Drosophila* is attracted to or repelled from the activation of specific homotypic uPNs. For example, DA2 responds to geosmin, a chemical generated from harmful bacteria and mold, which evokes a strong repulsion in *Drosophila* (***Stensmyr et al., 2012***). Similarly, VM3 is suggested to encode repulsive odors, while VM2 and VM7d encode attractive odors (***Mansourian and Stensmyr, 2015***; ***Bates et al., 2020***). Overall, the following information is acquired from the literature (***Hallem et al., 2004***; ***Galizia and Sachse, 2010***; ***Mansourian and Stensmyr, 2015***; ***Badel et al., 2016***; ***Bates et al., 2020***) and labeled accordingly:

- DA1, DA3, DL3, DM1, DM4, VA1v, VA2, VA3, VC1, VC2, VM1, VM2, VM4, VM5d, VM5v, VM7d, and VM7v (17 homotypes) encode attractive (O) odor.
- D, DA2, DA4l, DA4m, DC1, DC2, DC3, DC4, DL1, DL4, DL5, DM2, DM3, DM5, DM6, DP1m, V, VA5, VA6, VA7l, VA7m, VC3, VL2a, VL2p, and VM3 (25 homotypes) encode aversive (×) odor.
- The remaining homotypes are characterized as either unknown, non-preferential, or conflicting valence information.

Collecting the glomerular types of tightly bundled homotypic uPNs with *λ*_*X*_ < 0.4 in LH (Figures 6A and 7), we explore the presence of any organizational trend.

1. In LH, out of 37 homotypes composed of multiple uPNs based on our selection criterion (2 ≤*n* ≤8), 29 glomeruli (DL2v, DL2d, VM1, VL1, DM6, VM7d, VA3, VM5v, DA3, VM2, DL1, VA7m, VC3, VM7v, VC4, V, DM2, VM3, DA2, D, DC2, VA5, VA1v, DA1, DC3, DL3, VA1d, VP1d, and VP1l) satisfy the condition of *λ*_*X*_ < 0.4.
2. Homotypes VA1v, DA1, DC3, DL3, and VA1d (colored purple in Figures 3, 4, 6A, and 7) encode pheromones involved with reproduction (***Grabe et al., 2016***; ***Bates et al., 2020***; ***Dweck et al., 2015***), and VM4, VM1, VM7d, DM1, DM4, VC2, VM5d, VA3, VM5v, DA3, and VM2 encode odors presumed to be associated with identifying attractive food sources (***Couto et al., 2005***; ***Semmelhack and Wang, 2009***; ***Mohamed et al., 2019***; ***Bates et al., 2020***) (see Figure 6A). A previous work (***Grosjean et al., 2011***) has identified a group of glomeruli that co-process food stimuli and pheromones via olfactory receptor gene knock-in coupled with behavioral studies. The list of homotypes mentioned above is largely consistent with those glomeruli reported by ***Grosjean et al***. (***2011***).
3. Homotypes DM6, DM2, VM3, VL2p, DA2, and D are likely associated with aversive food odors. DA2 responds to bacterial growth/spoilage; VL2p, DM2, and VM3 to the alcoholic fermentation process; DM6 and D to flowers (***Galizia and Sachse, 2010***; ***Bates et al., 2020***).
4. Many homotypes responding to odors which can be described as kairomones, a type of odors emitted by other organisms (***Kohl et al., 2015***), are part of the 29 homotypes with *λ*_*X*_ < 0.4. This includes the pheromone encoding groups (VA1v, DA1, DC3, DL3, and VA1d) and others such as DA2, VC3, and VA5, which respond to geosmin, 1-hexanol, and 2-methyl phenol, respectively (***Hallem et al., 2004***; ***Galizia and Sachse, 2010***).

Figure 8 recapitulates the cluster information from *d*_*αβ*_-based analysis along with homotypes, odor types (color-codes), and putative valence (attractive (O) and aversive (×) odors). A few points are worth making:

**Figure 8.**
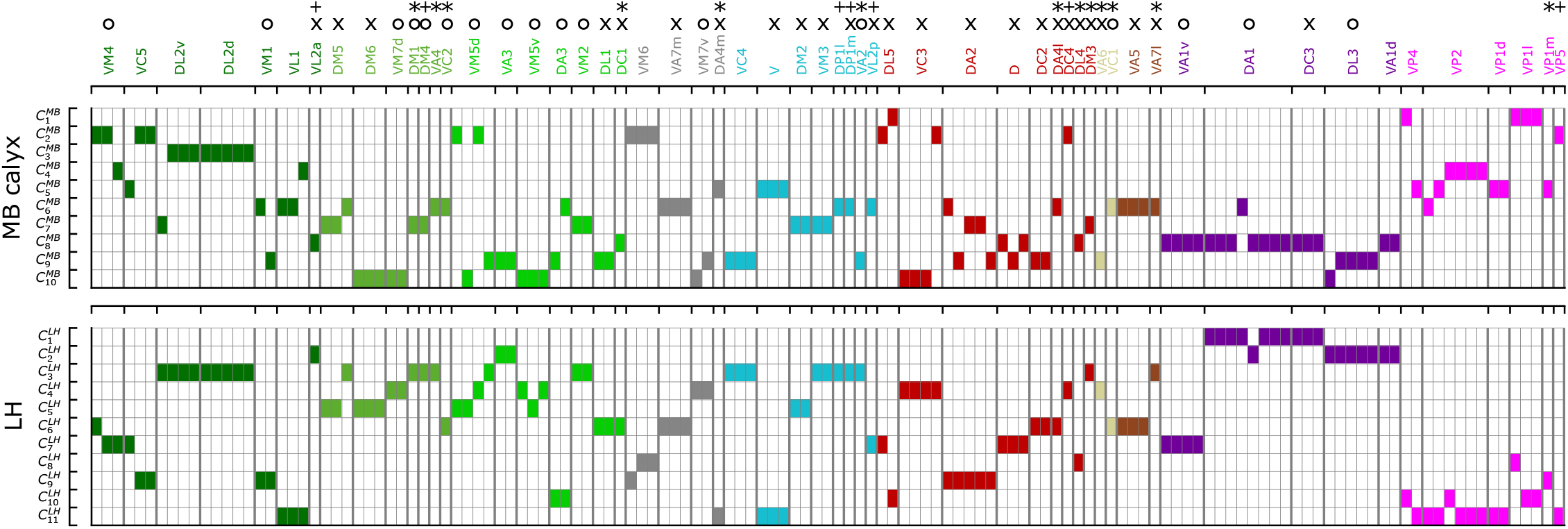
A diagram summarizing how the clusters of uPNs in MB calyx (10 clusters) and LH (11 clusters) are associated with the odor types (Dark green: decaying fruit, olive: yeasty, green: fruity, cyan: alcoholic fermentation, red: general bad/unclassified aversive, beige: plant matter, brown: animal matter, purple: pheromones, gray: unknown, pink: hygro/thermo). Asterisks (*) mark homotypes composed of a single uPN while plus (+) mark homotypes composed of a single uPN under our selection criterion but are actually a multi-uPN homotype, whose intra-homotype uPN distance is not available. O and × represent the putative valence information collected from the literature (O: attractive, ×: aversive).

1. Even though uPNs innervating MB calyx exhibit large *λ*_*X*_ s, the hierarchical clustering grouped homotypic uPNs together. This suggests the homotypic uPNs are still proximal in MB calyx, indicating the reduction in *d*_inter_ is what is driving the increase in overlapping. This is already shown through 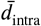 in Figure 5A and is supported by our statistical tests (see Tables S1 and S2). The same is true for LH.
2. 13 out of 57 glomeruli are made of a single uPN (*n* = 1, the asterisked glomeruli in Figures 6A and 8), which tend to be characterized by comparatively dense branched structures (see Figure S5), suggestive of homotypic uPN number dependence for the neuron morphology. Among the 13 homotypes, 7 encode aversive stimuli (×), 4 encode attractive stimuli (O), and 2 have no known valence information (see Table S3). The relative prevalence of single-uPN homotypes encoding aversive stimuli is noteworthy.
3. In LH, the cluster 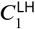, located in the anterior-ventral region of the neuropil, is composed only of pheromone-encoding homotypic uPNs, DA1 and DC3. The cluster 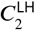 is also mostly composed of pheromone-encoding homotypic uPNs, DL3 and VA1d (Figures 4 and 8), which is consistent with the results by ***Jefferis et al***. (***2007***). In MB calyx, the majority of the uPNs encoding pheromones, except DL3, are grouped into the cluster 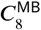 (see Figures 3 and 8).
4. Hygro/thermo-sensing homotypes such as VP2 and VP4 are spatially segregated from other odor-encoding uPNs. In MB calyx, these neurons rarely project ventrally and are distributed along the base of the neuropil. In LH, they are clustered in the dorsal-ventral-medial region, hardly innervating the neuropil but covering the medial side of the neuropil (Figures 3 and 4).
5. Along with the clusters of uPNs visualized in Figures 3 and 4, of particular note are the clusters formed by a combination of several homotypic uPNs. A large portion of uPNs innervating LH that encodes potentially aversive responses are grouped into clusters 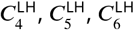, and 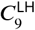, which envelop the cluster 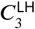 where mostly food-related homotypes converge (Figure 4).

Given that the synaptic communications established with KCs and LHNs are critical for neural computation in the inner brain, the specific type of uPN organization in each neuropil should be of great relevance. Indeed, it has been suggested that the spatial convergence, segregation, and overlapping of different homotypic uPNs within neuropil influence the information processing in higher olfactory centers (***Grosjean et al., 2011***).

According to previous studies (***Jefferis et al., 2007***; ***Liang et al., 2013***; ***Kohl et al., 2013***; ***Fisek and Wilson, 2014***), uPN innervation in LH and LHNs are highly stereotyped in terms of connectivity and response. Homotypic uPNs are spatially organized in AL, and to a certain degree, in LH, based on the odor type and valence information (***Min et al., 2013***; ***Huoviala et al., 2020***). The presence of tightly bundled anatomy of homotypic uPNs (*λ*_*X*_ < 0.4) in both AL and LH (Figure 7B) may imply that the *Drosophila* olfactory system dedicates a part of the second-order neural circuit on behalf of the *“*labeled-line” design, which enables the organism to sense urgent chemical stimuli at the early stage of information processing without going through more sophisticated neural computation in the inner brain (***Howard and Gottfried, 2014***; ***Andersson et al., 2015***; ***Min et al., 2013***).

### Labeled-line design of the higher order olfactory neurons

The concept of labeled-line design is widely considered at work at the ORN-PN interface (AL) as the signal generated from specific olfactory receptors converges to a single glomerulus (***Vosshall et al., 2000***; ***Couto et al., 2005***; ***Fishilevich and Vosshall, 2005***). However, it has been suggested that potential labeled-line strategy or separated olfactory processing of aversive odors encoded by DA2 (***Stensmyr et al., 2012***; ***Seki et al., 2017***) and pheromone-encoding homotypes in LH (***Jefferis et al., 2007***; ***Ruta et al., 2010***; ***Kohl et al., 2013***; ***Frechter et al., 2019***; ***Bates et al., 2020***; ***Chakraborty and Sachse, 2021***) are also at work in specific third-order olfactory neurons. So far, we have shown that the labeled-line design is present in the higher olfactory centers of second-order neurons such as MB calyx and LH, where homotypic PNs are tightly bundled together despite the lack of glomerular structure. In this section, we will conduct a comprehensive analysis of the synaptic connectivity between PNs and third-order olfactory neurons (KCs and LHNs) using three demonstrations. We ask whether the labeled-line strategy implied in the PN organization is translated over to the third-order olfactory neurons, to what extent the signals encoded by different homotypic uPNs are integrated at synaptic interfaces with the third-order neurons, and whether the spatial properties of pre-synaptic neurons (PNs) play any role in signal integration by the third-order neurons.

### Homotype-specific connections

For the analysis of the interface between homotypic uPNs and third-order neurons, we study the connectivity matrices *C*^PN−KC^ and *C*^PN−LHN^ (see Figures 9A and S6), which are extracted from the hemibrain dataset (***Scheffer et al., 2020***). The *C*^*ξ*^ (*ξ* = PN-KC or PN-LHN) is a binary matrix (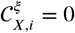 or 1 dictating the connectivity) of synaptic connectivity between *X*-th homotypic uPNs and *i*-th third-order neuron (KC or LHN).

**Figure 9.**
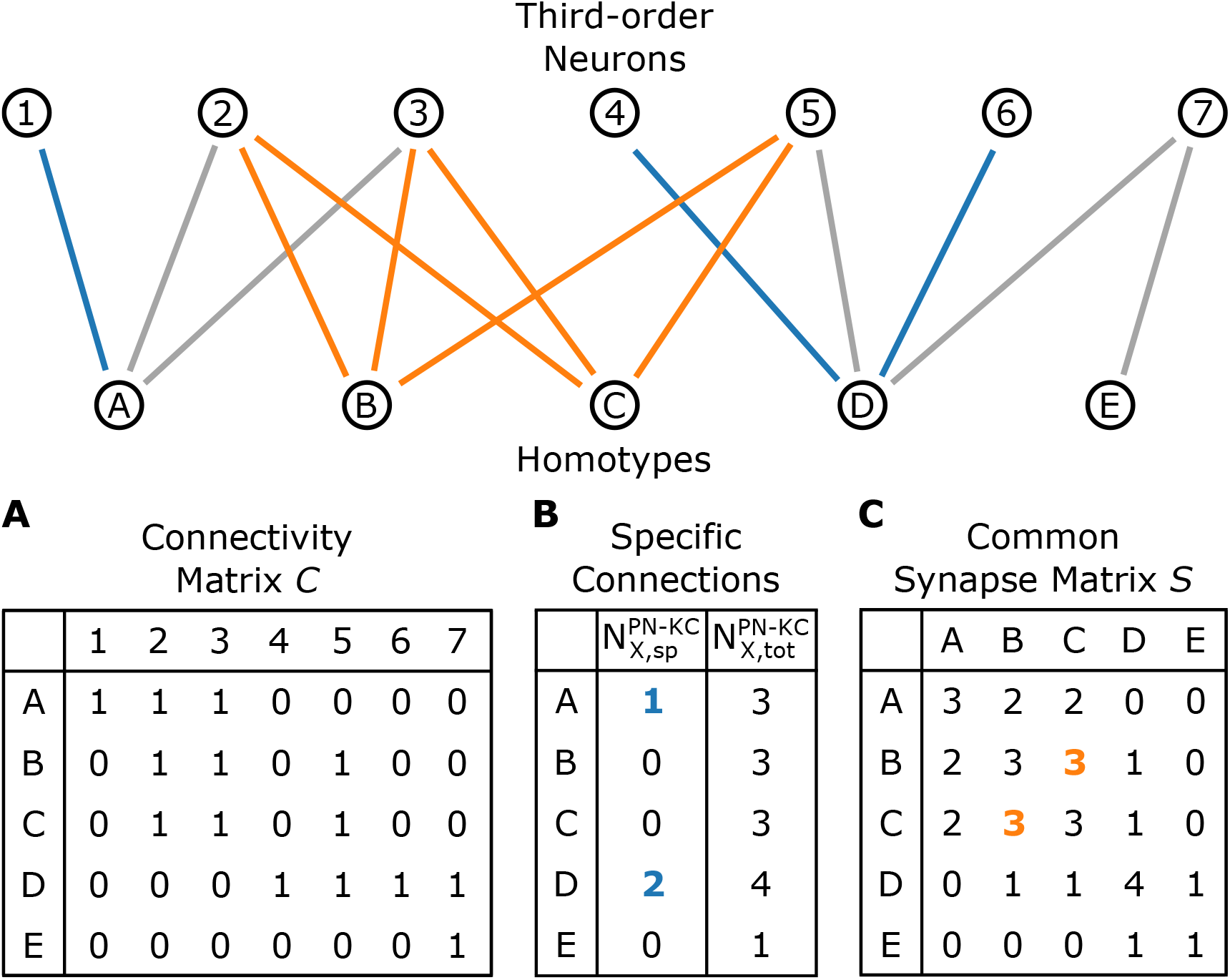
A schematic illustrating the connectivity between homotypes (*X* = *A, B*, …, *E*) and third-order neurons (*i* = 1, 2, …, 7). (A) The connectivity matrix *C. C*_*X,i*_ = 1 when any uPNs in the *X*-th homotype and *i*-th third-order neuron synapses and *C*_*X,i*_ = 0 otherwise. (B) The number of *X*-th homotype-specific connections (*N*_*X*,sp_) and the total number of third-order neurons synapsed to any uPNs in the *X*-th homotype. (C) The common synapse matrix (*S*) whose element speci1es the number of third-order neurons commonly connected between two homotypes. The homotype A is connected to three third-order neurons 1, 2, and 3 (*N*_*A*,tot_ = 3). Neuron 1 is not synapsing with any other homotype but A, and hence *N*_*A*,sp_ = 1; similarly, *N*_*D*,sp_ = 2 (the blue lines depict specific connections). The signals from the two homotypes B and C are shared by the third-order neurons 2, 3, and 5; therefore, *S*_*BC*_ = 3 in the common synapse matrix *S*.

The total number of third-order neurons in contact with *X*-th homotypic uPNs can be obtained by counting the non-zero elements of the matrix *C* with fixed *X* (For the case of PN-KC interface, this number can be obtained from 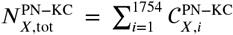). Among the third-order neurons in contact with *X*-th homotypic uPNs, a group of third-order neurons that synapse solely with *X*-th homotypic uPNs can be identified (see the explanation in Figure 9B). For example, Figure 10A shows 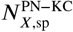, which denotes the number of *X*-th PN specific-KCs, and those normalized by 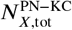 (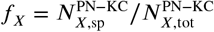, see Methods for the detailed algorithms behind the calculation), for all homotypes (*X* = VM4, VC5, …, VP4). Compared to those in KCs, the *‘*homotype-specific*’* connections are much more prevalent in LHNs (Figure 10). Certain homotypic uPNs, e.g., hygro/thermo-sensing homotypes are connected to the LHNs which are dedicated to process the signals from hygro/thermo-sensing homotypes (≥10% of PN-LHN connections made by homotypes). The existence of these *‘*homotype-specific*’* third-order neurons suggests that a subset of olfactory processing may rely on the labeled-line strategy that extends beyond the layer of second-order neurons to the higher brain center.

**Figure 10.**
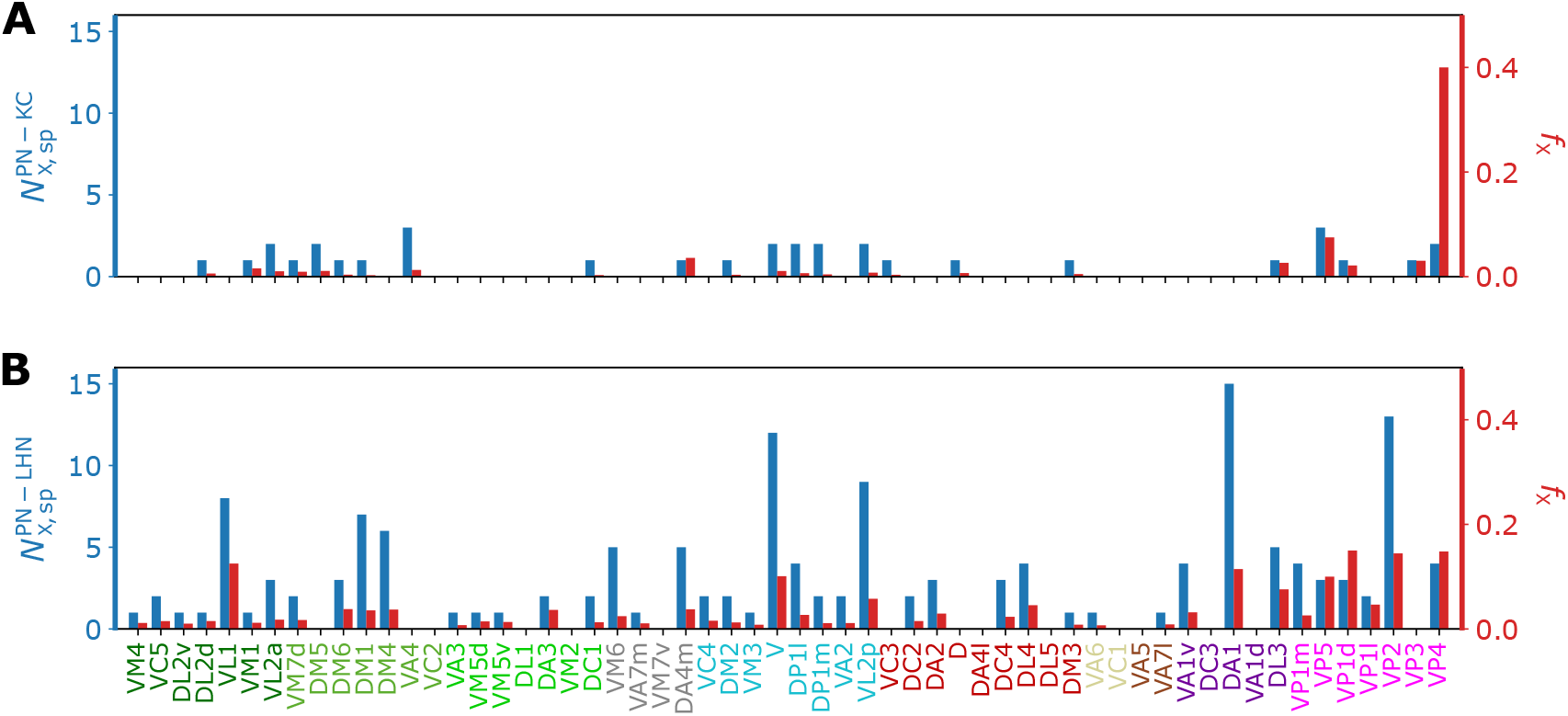
Bar graphs depicting the number of KCs/LHNs that synapse with a specific homotype *X* (*N*_*X*,sp_, blue) and the percentage of KCs/LHNs that synapse with a specific homotype *X* (*f*_*X*_ = *N*_*X*,sp_/*N*_*X*,tot_, red) at (A) PN-KC and (B) PN-LHN interfaces.

### Third-order neuron mediated signal integration

Figures 11A and 11B show the *‘*common synapse matrices*’* representing the number of commonly connected third-order neurons between two homotypes *X* and *Y* (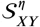 with *η* = MB or LH), which provide glimpses into the extent of signal integration mediated by KCs and LHNs (see Figure 9C and the caption for how these matrices are constructed from the connectivity matrix).

**Figure 11.**
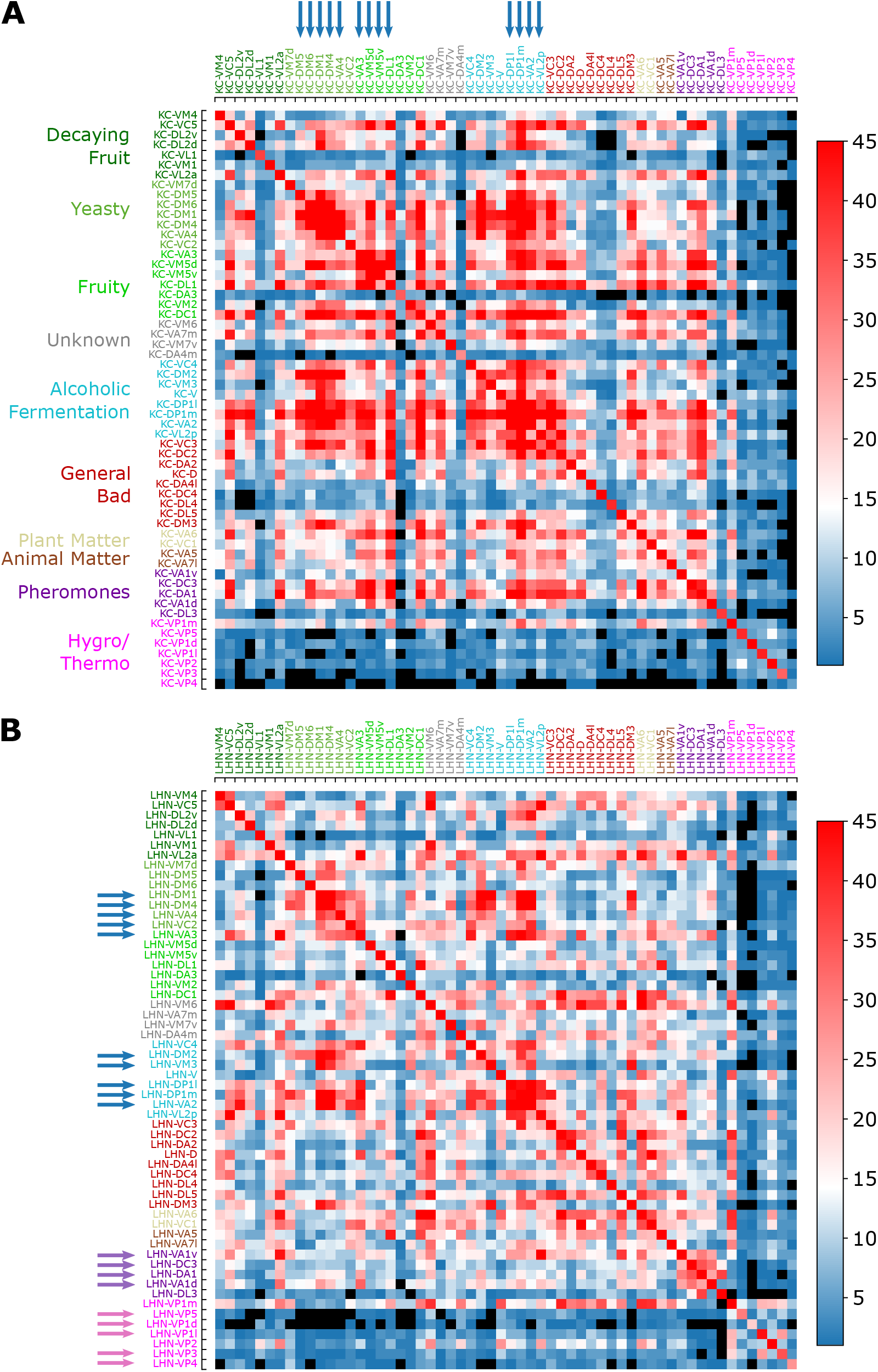
Common synapse matrices (A) *S*^KC^ and (B) *S*^LHN^, each of which represents the extent of signal integration from homotypic uPNs to KCs and LHNs. The black color is used when there is no third-order neuron-mediated signal integration (*S*_*XY*_ = 0) happening between two homotypes *X* and *Y*. See Figure 9C and its caption for how the common synapse matrices are calculated from the connectivity matrices provided in Figure S6.

1. Overall, the number of synaptic connections between uPNs and KCs is greater than that between uPNs and LHNs (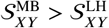, see Figure S7).
2. 2.In MB calyx, the signals from food-related odors-encoding homotypes (e.g., Yeasty, Fruity, or Alcoholic Fermentation odor types) are shared by a large number of KCs, which constitute a few large clusters in *S*^MB^ matrix, depicted in red 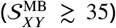 and indicated by the blue arrows on the top in the Figure 11A). Some KCs process signals almost exclusively from the hygro/thermo-sensing homotypes without sharing any signal from other homotypes (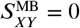 for the cases of *X* and *Y* homotype pairs without any signal integration, which are depicted in black in Figure 11). There are also homotypes with significantly less number of overall synaptic connections to KCs, dictated by the diagonal element of the matrix *S*^MB^ (see Figure S7A). In comparison with *S*^LH^, the *S*^MB^ suggests a stronger but less organized signal integration between heterotypic PNs by KCs and lends support to the previous literature pointing to the random synapsing of KCs with uPNs at MB calyx (***Caron et al., 2013***; ***Stevens, 2015***; ***Eichler et al., 2017***; ***Zheng et al., 2020***).
3. *S*^LH^, on the other hand, demonstrates LHN-mediated signal integration localized to subsets of homotypes. First, the pheromone-encoding and hygro/thermo-sensing homotypes share the synaptic connections to LHNs among themselves, which are demonstrated as the homotype-specific block patterns along the diagonal of *S*^LH^ matrix (see purple and pink arrows on the side in Figure 11B). The *S*^LH^ matrix also shows that signals from various food-related odor encoding homotypes, such as DP1l, DP1m, VA2, and VL2p or DM1, DM4, and VA4 are also integrated (see blue arrows in Figure 11B). Many of these homotypes encode signals originating from esters, which is intriguing given the ester-encoding LHN cluster shown by ***Frechter et al***. (***2019***). The results suggest that certain odor types are processed through common channels of LHNs that are largely dedicated to encoding a particular odor type.

### Spatial proximity-based versus connectivity-based clustering

Next, we study the relationship between spatial proximity-based clustering and connectivity-based clustering results. Upon visual inspection, the connectivity-based clustering at MB calyx (Figure 12A on the right) appears less structured than the spatial proximity-based clustering (Figure 12A on the left). For example, many homotypic uPNs are grouped under a common branch in the tree structure obtained from the spatial proximity-based clustering, whereas such a feature is largely absent in the output of the connectivity-based clustering. Therefore, the spatially well-clustered uPNs at MB calyx do not precisely translate to structured connectivity patterns, consistent with the notion of randomized PN-KC connections (***Caron et al., 2013***; ***Stevens, 2015***; ***Eichler et al., 2017***; ***Zheng et al., 2020***). In stark contrast to the outcomes for MB calyx, most homotypic uPNs are grouped in the connectivity-based clustering for LH (Figure 12B). This suggests that the spatially proximal uPNs synapse with a similar group of LHNs. The stereotyped organization and connectivity of uPNs in LH have been suggested before (***Jefferis et al., 2007***; ***Liang et al., 2013***; ***Kohl et al., 2013***; ***Fişek and Wilson, 2014***), and we demonstrate such stereotypy is, in reality, expressed throughout LH over all uPNs. In LH, spatial and organizational characteristics of uPNs are well-translated to connectivity to LHNs. A quantitative comparison of two trees based on statistical tests lends support to the notion that the spatial organization of uPNs is indicative of connective properties, most evident in LH (see Supplementary Information for Baker’s Gamma index, entanglement, and cophenetic distance correlation).

**Figure 12.**
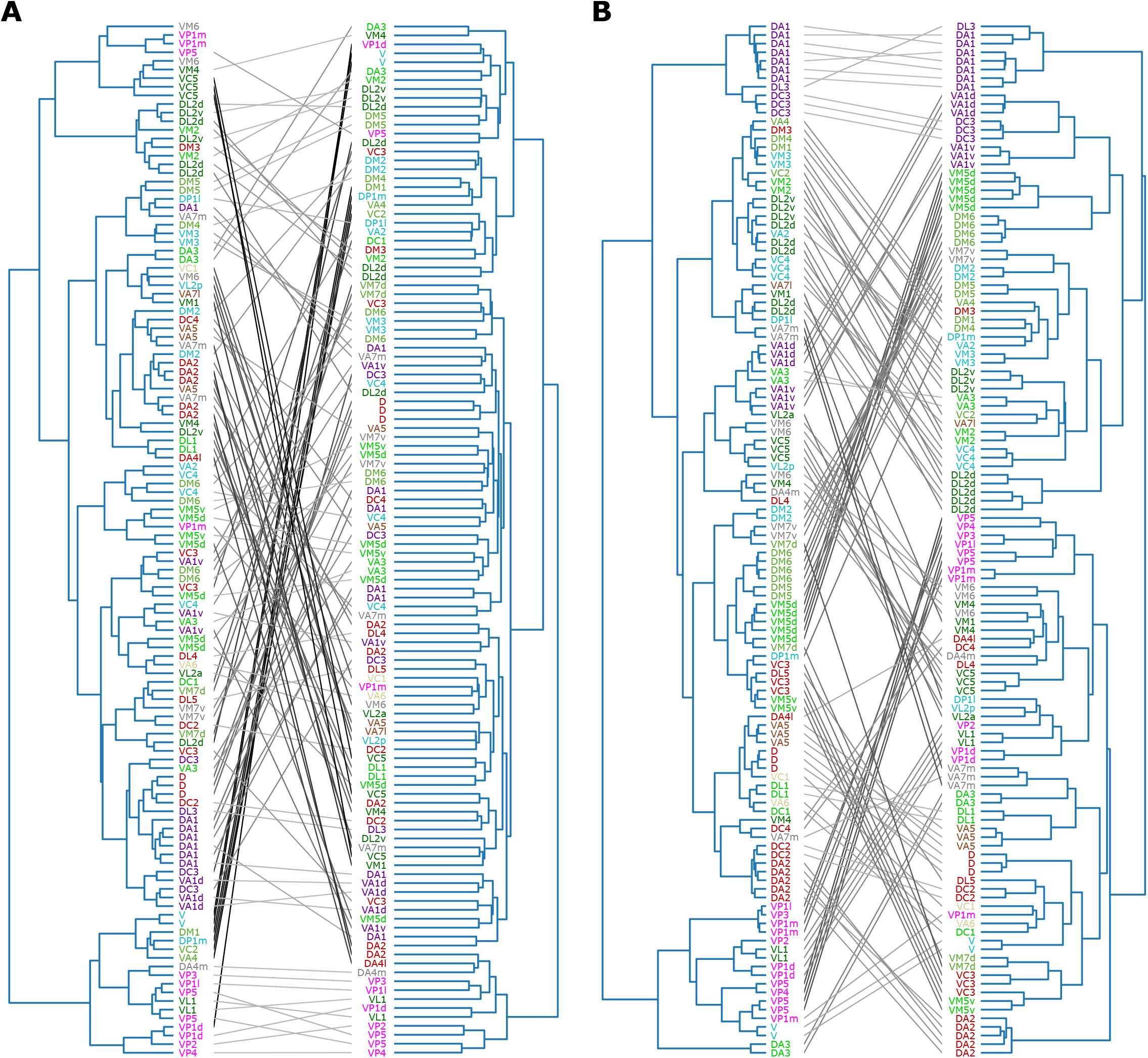
Tanglegrams comparing the tree structures generated from the inter-PN distances-based (left) and the connectivity-based clustering (right) (A) between uPNs and KCs, and (B) between uPNs and LHNs. The same uPNs in the two tree structures are connected with lines, which visualize where the uPNs clustered by one method end up in the clustering results of another. The labels for uPNs are representative of the homotype and are color-coded based on the encoded odor types (Dark green: decaying fruit, lime: yeasty, green: fruity, gray: unknown/mixed, cyan: alcoholic fermentation, red: general bad/unclassified aversive, beige: plant matter, brown: animal matter, purple: pheromones, pink: hygro/thermo).

## Discussions

The inter-PN organization revealed in this study and its association with odor type/valence are reminiscent of the generally accepted notion that *the form determines the function* in biology. Previously observed stereotypes of neurons in the *Drosophila* olfactory system were largely based on the differentiation between pheromones and non-pheromones (***Ruta et al., 2010***; ***Kohl et al., 2013***; ***Frechter et al., 2019***; ***Chakraborty and Sachse, 2021***), the whole-cell patch-clamp recording (***Seki et al., 2017***), and imaging studies suggestive of stimulus-dependent arrangement of neurons in LH (***Marin et al., 2002***; ***Wong et al., 2002***; ***Jefferis et al., 2007***). Our results are generally consistent with the previous studies, which suggest that a level of stereotypy in uPN organization in MB calyx and LH is universal throughout *Drosophila*, which can be captured through different metrics and methodologies. In line with ***Lin et al***. (***2007***), our study finds that homotypes DL2v and DL2d constitute a bilateral cluster in MB calyx 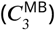, and that the dual organization of uPNs is present in MB calyx and LH, such that homotypes DC2, DL1, and VA5 are sorted into the same cluster in LH while sharing similar innervation pattern in MB calyx. Our clustering results in LH share similarities with the NBLAST score-based LH clusters (***Bates et al., 2020***). The uPNs that ended up in the same cluster or nearby clusters, such as homotypes DM1, DM3, DM4, VA4, and VM3 in the cluster 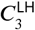, are also grouped in the NBLAST score-based clustering analysis (***Bates et al., 2020***). We find a significant correlation of *d*_*αβ*_ with NBLAST score (see Figure S1) despite the fact that two metrics prioritize different aspects of neuronal anatomy.

Our inter-PN distances and clustering results suggest the spatial organization of uPNs differs greatly in each neuropil (Figure 5). Some of the tightly bundled organization of PN homotypes are well preserved throughout the neuropils despite the lack of glomerulus in MB calyx and LH. The spatial segregation between different homotypes is, however, practically not present in MB calyx, leading to a high degree of overlapping. Therefore, the heterogeneity of homotypes at the PN-KC synaptic interface may physically assist the randomized sampling known to exist between PNs and KCs (***Caron et al., 2013***; ***Stevens, 2015***; ***Eichler et al., 2017***; ***Zheng et al., 2020***).

Our analysis suggests that LH is compartmentalized into four regions: (1) Dorsal-posterior region primarily occupied by food-related uPNs; (2) Ventral-anterior region occupied by pheromone-encoding uPNs; (3) Biforked bundle surrounding dorsal-posterior region largely occupied by food-related uPNs with an aversive response; (4) Dorsal-anterior-medial region occupied by hygro/thermo-sensing uPNs. Previous attempts at identifying regions of odorant space in LH revealed compatible results. The three domains (LH-PM, LH-AM, and LH-AL) suggested by ***Strutz et al***. (***2014***) seem to be a different combination of our clustering result (LH-PM and LH-AM correspond to the dorsal-posterior region and LH-AL corresponds to a combination of ventral-anterior region and the biforked bundle). Although not perfect, the study of the axo-axonic communities in LH yields results with comparable characteristics (***Bates et al., 2020***), understandably due to the necessity of inter-neuronal proximity to form synapses. For example, community 12 by ***Bates et al***. (***2020***) is predominantly composed of homotypes VP1l and DL5, which resembles our cluster 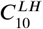. Community 6 contains a mixture of homotypes VA5, VC1, D, DA4l, DC2, DA3, and VA7m, which is reminiscent of our cluster 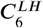.

Many homotypic uPNs that are spatially localized in LH can be associated with key survival features and a strong innate response (***Seki et al., 2017***). In this sense, the stereotyped localization of pheromone-encoding uPNs in 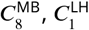, and 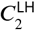 is of great interest. Our study not only lends support to the existing studies pointing to the labeled-line strategy in the *Drosophila* olfactory system but also suggests that an even more sophisticated level of spatial organization, which supersedes the pheromone versus non-pheromone segregation, may exist. Interestingly, while the spatial organization of uPNs in LH has a basis on the functionality of the odor type encoded, it does not seem to be directly translated to segregated chemical features seen in LHNs (***Frechter et al., 2019***). The apparent divergence observed at the PN-LHN interface, coupled with strongly stereotyped connectivity may contribute to a higher resolution of odor categorization.

Our study suggests that while the primary connectivity motif of third-order olfactory neurons indeed integrates signals, there still exist several labeled lines. The synaptic connections at the PN-KC interface are largely integrative and randomized - with an exception of hygro/thermo-sensing homotypes. The uPNs in LH are spatially segregated, which translates to connectivity in three different levels. First, certain LHNs are dedicated to encoding signals from a specific homotype. The number of these *‘*homotype-specific*’* LHNs varies across the homotype and can make up a significant portion of PN-LHN connections depending on the homotype (Figure 10). Second, synaptic connectivity maps between uPNs and LHNs indicate odor type-dependent integration occurs at LH (Figure 11B). Channels of LHNs predominantly encoding specific odor types are observed; one primarily integrates responses from certain food-related homotypes, one integrates pheromone-encoding homotypes, and another integrates hygro/thermo-sensing homotypes. Third, homotypic uPNs share similar connectivity to LHNs, unlike those in MB calyx. The signals relayed from the spatially well-organized (or tightly bundled) homotypes are localized into a specific group of LHNs, thereby forming a *‘*homotype-specific*’* connectivity motif (Figures 10, 11, and 12).

In our study of the labeled-line strategy, we made several interesting observations, which are worth comparing with the concept of *‘*fovea*’* introduced by ***Zheng et al***. (***2020***). A *‘*fovea*’* delineates deviations between experimentally observed connectivity matrices and connectivity under the assumption of random synapses in MB calyx, specifically for certain food-related uPNs (***Zheng et al., 2020***). A group of common KCs predominantly sampling *‘*food-related*’* uPNs manifest themselves in the common synapse matrix *S*^KC^ (see the group of homotypes comprising the clusters, highlighted by the blue arrows in Figure 11A). A subset of homotypic uPNs under the food-related *‘*fovea*’* reported by ***Zheng et al***. (***2020***) are also spatially clustered (e.g. DM1, DM4, DP1m, DP1l, VA2, and VA4). While most of these homotypes are spatially proximal (the vast majority of the uPNs are located in clusters 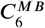 and 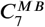), some homotypes under the food-related *‘*fovea*’* such as VA2 are sampled from spatially disparate clusters. Thus, it is likely that factors other than the spatial organization of uPNs in neuropils contribute to creating the *‘*fovea*’*. Interestingly, the spatial proximity of pheromone-encoding homotypes in MB calyx may suggest the existence of pheromone-encoding *‘*fovea,*’* but most uPNs in these homotypes do not converge in connectivity-based clustering with an exception of VA1d. In fact, we suspect the spatial organization of pheromone-encoding homotypes in MB calyx, which is placed at the center of the neuropil, to facilitate the observed randomization of connections by increasing the accessibility of KCs to these homotypes. There is, however, a potential hygro/thermo *‘*fovea,*’* where homotypes such as VP1d and VP2 are spatially clustered together and the signals from these homotypes are relayed by the same set of KCs. Curiously, VL1 is part of this hygro/thermo *‘*fovea*’* (Figure 12A).

To show that the spatial and organizational properties of uPNs we observed for the FAFB dataset are general, we have analyzed the hemibrain dataset (***Scheffer et al., 2020***) and carried out the same calculation to generate some of the main figures in our study (see Figure S8 and Supplementary Information). Remarkably, the results from the hemibrain dataset are consistent with those from the FAFB dataset. For example, 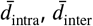, and *λ* are almost identical between both datasets (see Figures 5 and S8A). 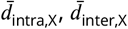, and *λ*_*X*_ show slight differences due to a mismatch between the FAFB and the hemibrain dataset (on glomerulus labels and the number of uPNs based on our selection criterion) leading to a different number of uPNs per homotype (Figure S8B), but the correlation between *λ*_*X*_ s at MB calyx and LH are still observed (Figure S8C). Most importantly, the clustering results are similar, where many spatial clusters in both datasets share the same set of homotypes. Additionally, odor type-dependent spatial properties are retained (Figure S8D), with all statistical tests supporting our hypothesis. In conclusion, our observations seem generalizable, lending support to the previous claims of stereotypy in the *Drosophila* brain and neuronal structures (***Jenett et al., 2012***; ***Jeanne et al., 2018***; ***Schlegel et al., 2021***).

Apart from uPNs primarily explored in this study, a host of local neurons (LNs) and multiglomerular PNs (mPNs) also constitute sophisticated neural circuits to regulate the signals received from ORNs (***Sudhakaran et al., 2012***; ***Bates et al., 2020***), playing a significant role in the olfactory signal processing (***Olsen et al., 2010***; ***Jeanne and Wilson, 2015***; ***Seki et al., 2017***). A large portion of these mPNs is GABAergic and inhibitory (***Berck et al., 2016***; ***Tobin et al., 2017***), whereas the role of interneurons can be both inhibitory and excitatory (***Turner et al., 2008***; ***Wilson et al., 2004***). Electrophysiological measurements indicate that mPNs are narrowly tuned to a specific set of odor stimuli (***Berck et al., 2016***), which is significant given that PNs are generally thought to be more broadly tuned than presynaptic ORNs (***Wilson et al., 2004***). Several PNs do not follow the typical mALT, but mediolateral antennal lobe (mlALT) or lateral antennal lobe tracts (lALT) instead, thereby bypassing innervation through one of the higher olfactory centers (***Schultzhaus et al., 2017***; ***Zheng et al., 2018***; ***Bates et al., 2020***). As stated previously, we confined ourselves to uPNs innervating all three neuropils to compare the spatial organization across neuropils for each uPN. As a result, 28 uPNs present in the FAFB dataset are not explored in our study. In MB calyx, only two uPNs constituting VP3 were dropped, which ended up in an almost identical clustering output once hierarchical clustering was performed on the entire 137 uPNs that innervate MB calyx. Two missing uPNs were grouped into clusters 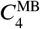 and 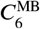, along with other hygro/thermo-sensing homotypes. On the other hand, the addition of 27 uPNs constituting 15 homotypes innervating LH but not MB calyx created four new clusters when hierarchical clustering was performed (Figure S9). The additional uPNs changed the content of the individual clusters; that is, the tree-cutting algorithm broke down a few clusters that became larger due to the additional uPNs. Furthermore, when we calculate the 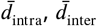, and *λ* in LH for the 15 homotypes that included the 27 uPNs, we find that the 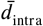 values increased when the 27 uPNs were included (see Figure S10). This suggests that the previously removed uPNs, most of which follow mlALT, are significantly different in terms of spatial and organizational characteristics and thus should be analyzed separately. Out of 27 additional uPNs in LH, 21 were in mlALT, 5 were in trans-lALT, and 1 was in mALT. Figure S11 illustrates how these 27 uPNs innervate LH which demonstrates the reason behind increased 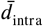 values. These 27 uPNs are mostly GABAergic (21 are labeled as GABAergic, 1 as cholinergic, and 4 as unknown neurotransmitter type), covering 84% of all GABAergic uPNs available in the FAFB dataset. These uPNs innervate LH differently from other uPNs in the same homotype that follow mALT (see homotypes such as DA1, DC4, DL2d, DL2v, DP1l, VA1d, VA1v, VL2a, VL2p, and VP5 in Figure S11). Morphologically, inhibitory GABAergic neurons are often considered *‘*smooth*’* and aspiny (***Douglas et al., 1989***; ***Bopp et al., 2014***; ***Gouwens et al., 2019***), which are discernible from Figure S11.

It is of great interest that many of the single-uPN homotypes, characterized by densely branched morphology, encode signals with aversive responses. Direct transmission of the associated signals across the three neuropils via a single PN might simplify the overall processing of the olfactory signals as well as reduce the energetic cost. Similarly, the morphological characteristics of uPN innervation at each neuropil are intriguing. Even though a structural difference exists between the single-uPN and multi-uPN homotypes, all uPN innervations within neuropil share a similar mophology regardless of the homotype (see Figure S5) (***Choi et al., 2022***). A localized morphological diversity within a neuron may be a characteristic aspect of pseudo-unipolar neurons like uPN and suggests a fundamentally multi-scale characteristic of neuron morphology.

The *Drosophila* brain EM reconstruction project has evolved to its near completion since the EM image dataset was first released (***Dorkenwald et al., 2022***). The reconstruction of the majority of the *Drosophila* central brain as well as the corresponding connectome with detailed information of the individual synapses has become publicly available (***Scheffer et al., 2020***). Our analysis of the second-order neurons inside the *Drosophila* olfactory system may be translated to other parts of the nervous system in *Drosophila* as well as different organisms including the central nervous system (CNS) of humans. For the mammalian olfactory system, the details of analyses must be adapted, however, since the wiring scheme is much more complex than that of an insect (***Maresh et al., 2008***). For example, multiple glomeruli encoding the same olfactory signal exist in humans (***Mombaerts et al., 1996***). When analyzing the spatial properties, this can be accounted for by prioritizing the individual glomerulus over the homotypes. Then, homotypic PNs forming different glomeruli may be compared or averaged if one were to consider the homotype-dependent characteristics. According to the neurotransmitter map from a recent study (***Dolan et al., 2019***), sophisticated processes beyond neuronal anatomy are apparently at work in the olfactory signal processing. Thus, functional studies incorporating odor response pro*file*s in PNs (***Badel et al., 2016***) and ORNs (***Münch and Galizia, 2016***; ***Bak et al., 2018***) would supplement our findings. The extension of our study to the other regulatory interneurons and mPNs, morphological studies of second-order neurons, and spatial analysis of third-order neurons will be of great interest for a better understanding of the olfactory signal processing beyond the implication of the neural anatomy and connectivity studied here.

## Materials and methods

### Data preparation

We used the neuron morphology reconstruction of 346 *Drosophila* olfactory neurons from the FAFB dataset (***Bates et al., 2020***) traced from EM images. The neurons were extracted from the right hemisphere of the female *Drosophila*. Out of 346 neurons in the FAFB dataset, 164 neurons were uPNs. One uPN in the dataset (neuron ID = 1356477 forming VP3) did not have an associated reconstruction (.swc file) available and was therefore ignored. For this study, we chose uPNs that innervate all three neuropils because 1) we want to compare spatial characteristics of the uPN innervation across each neuropil and 2) to classify each uPN based on the odor encoding information. Out of 164 uPNs, we selected uPNs that innervate all three neuropils and collected a total of 135 uPNs constituting 57 homotypes under this criteria. This criterion resulted in mostly cholinergic uPNs that follow mALT. Rest of the uPNs that did not innervate all three neuropils are collected for the supplementary analysis. The morphological information of each neuron is stored as a set of 3D coordinates with the connectivity specified with the parent nodes. Complete reconstruction of neuron morphology was made by connecting data points based on their parent-child relationship.

For the reproducibility study, we used the hemibrain dataset (***Scheffer et al., 2020***) taken from the neuPrint database (***Clements et al., 2020***). We collected a total of 120 uPNs forming 58 glomeruli based on the same criterion we used for the FAFB dataset (uPNs that innervate all three neuropils) from the right hemisphere of the female *Drosophila*. Of the 120 uPNs from the hemibrain dataset, five uPNs had ambiguous glomerulus labels associated with them presumably due to poorly formed glomerular structures. For these uPNs, we took the glomerulus labels from the FAFB dataset with the matching hemibrain neuron IDs.

Additionally, a recent community-led effort identified three glomeruli with conflicting glomerulus labels which have been a source of confusion. The community agreed to update labels VC3l, VC3m, and VC5 to VC3, VC5, and VM6, respectively (***Schlegel et al., 2021***), which has been manually incorporated into our analyses for both the FAFB and the hemibrain dataset.

Next, we systematically demarcated the regions of AL, LH, and MB calyx. The density of data points projected to each axis was used for the identification since the neuropils are featured with a much higher density of data points than the rigid backbone connecting them. The boundaries defining each neuropil were systematically chosen from local minima that separate neuropils from rigid backbones. Due to the unique structure of uPNs, sometimes the projection along a given axis cannot fully differentiate two neuropils. To resolve this issue, projections along each axis were sampled while rotating the data points along the reference axes at 5° increments to obtain multiple snapshots. The densities were analyzed to choose the optimal degrees of rotation along the reference axes that could best segment the neuropils. We used the smallest average and deviation value of density at the local minima as the criteria to choose the optimal rotation. The process has been repeated for each neuropil to identify a set of boundaries along multiple transformed axes with various degrees of rotations that optimally confine each neuropil. This information has been combined to create a set of conditions per neuropil for segmentation. The resulting neuropils were confirmed through visual inspection. We compared our neuropil segmentation boundaries with neuropil volume surface coordinates provided by CATMAID (***Saalfeld et al., 2009***) and found the boundaries are comparable (data not shown). An overview of the segmentation process is available in Figure S12.

The odor type and odor valence information were extracted from various literature (***Hallem et al., 2004***; ***Galizia and Sachse, 2010***; ***Stensmyr et al., 2012***; ***Mansourian and Stensmyr, 2015***; ***Badel et al., 2016***; ***Bates et al., 2020***) and we closely followed the categorical convention established by ***Mansourian and Stensmyr*** (***2015***) and ***Bates et al***. (***2020***). However, we note that the categorization of a uPN under a specific odor category may overshadow the complete spectrum of odorants a uPN might encode, especially if the uPN encodes ORs that are broadly tuned. There-fore, we focused on the well-separated pheromone/non-pheromone encoding types and valence information.

To test our labeled-line hypothesis the connectivity information between uPNs and higher olfactory neurons such as KCs and LHNs was necessary. Since only the hemibrain dataset contains detailed connectivity information, all of our connectivity studies are done using uPNs, KCs, and LHNs queried from the hemibrain dataset. We chose KCs and LHNs with the synaptic weight greater than or equal to 3 for the 120 uPNs we tested in the reproducibility study. This resulted in 1754 KCs and 1295 LHNs, creating bipartite connectivity matrices at each neuropil.

### Inter-PN distance

The *“*distance*” d*_*αβ*_ between two neurons, *α* and *β*, with different lengths (*N*_*α*_ ≤ *N*_*β*_) is quantified by calculating

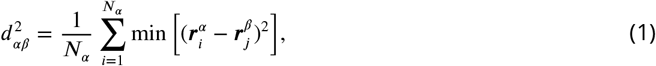

where 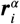 is an i-th coordinate forming the neuron *α*. Equation 1 is evaluated over all pairs of 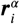 and 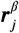 with *j* = 1, …, *N*_*β*_ that gives rise to the minimum value. This means that when *Nα ≤ N*_*β*_, for every i-th coordinate in the neuron 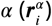, we find j-th coordinate in the neuron 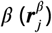that is the closest 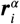. Then, the spatial proximity of a given pair of neurons is assessed by the *d*_*αβ*_ that denotes the average of all the minimum Euclidean distances between the pair of coordinates.

### The degree of bundling, packing, and overlapping

We define the mean intra- and inter-homotype distances as

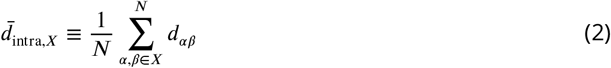

and

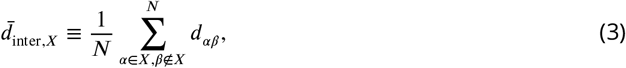

where *X* denotes a homotype and *N* is the total number of uPN pairs to be averaged. The 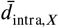 is calculated over all the pairs of uPNs in the *X*-th homotype, quantifying the tightness of bundling of uPNs constituting the *X*-th homotype. On the other hand, 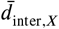 is calculated over the pairs of uPNs between *α* -th uPN belonging to the *X*-th homotype and *β* -th uPN in the *Y* -th homotype (*Y* ≠ *X*), such that it measures the extent of packing of uPNs around the *X*-th homotype. The degree of overlapping for the *X*-th homotype, *λ*_*X*_, is defined as the ratio of average intra- and inter-homotype distances,

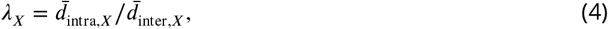

which represents how clearly the *X*-th homotype is segregated from other homotypes in a given space. If the value of *λ*_*X*_ is large (*λ*_*X*_ *»* 1), it implies that the space spanned by the *X*-th homotype is not clearly discerned from other homotypic uPNs.

### Spatial clustering of projection neurons

Hierarchical/agglomerative clustering was used to cluster the uPN innervation at each neuropil using the pairwise *d*_*αβ*_ matrices. First, the linkage was constructed by using the pairwise distance matrix using the Farthest Point Algorithm (or “complete” method), where the maximum distance between neurons is used to define the distance between two clusters. This criterion is used to build hierarchical relations (or nested clusters) in a bottom-up approach where each neuron is treated as a cluster at the beginning. The result is a fixed tree structure of individual neurons from which the finalized clusters are formed using an optimal tree-cutting algorithm. Looking at the dendrogram from AL (Figure S3A), homotypic uPNs are grouped together with high accuracy, suggesting our distance metric *d*_*αβ*_ is adequate. We tested various tree-cutting criteria such as elbow method, gap statistics, maximum average silhouette coefficient, and dynamic hybrid cut tree method (***Langfelder et al., 2008***) to determine the optimal number of clusters. We decided to use a method that gives a cluster number closest to the number of different odor types (which is 10). The dynamic hybrid cut tree method performed the best in this regard (Table 1). Therefore, we deployed the dynamic hybrid cut tree method with the minimum cluster size of four neurons for the tree-cutting, following the neuron clustering procedure used by ***Gouwens et al***. (***2019***).

**Table 1.**
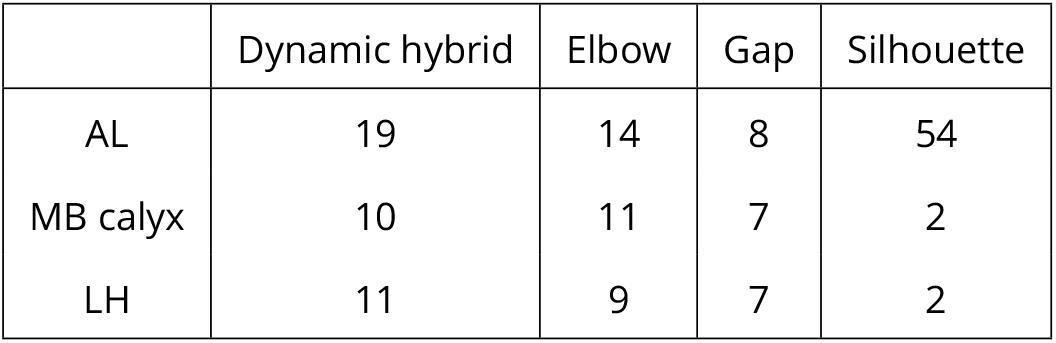
The optimal number of clusters according to the dynamic hybrid cut tree method, elbow method, gap statistics, and maximum average silhouette coefficient.

|  | Dynamic hybrid | Elbow | Gap | Silhouette |
| --- | --- | --- | --- | --- |
| AL | 19 | 14 | 8 | 54 |
| MB calyx | 10 | 11 | 7 | 2 |
| LH | 11 | 9 | 7 | 2 |

### Pearson’s *χ*^2^-test of independence

The association between two categorical variables is assessed using Pearson’s *χ*^2^-test. For the test, a contingency table, which lists the categorical frequency of two variables, is created. For example, *O*_*ij*_ of the *i*- and *j*-th element of the contingency table shown below is the frequency counting the putative valence *i* = 1 (attractive), 2 (aversive), 3 (unknown), and the number of uPNs in one of the 10 clusters in MB calyx with 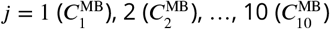.

| | $C_1^{\text{MB}}$ | $C_2^{\text{MB}}$ | $C_3^{\text{MB}}$ | $C_4^{\text{MB}}$ | $C_5^{\text{MB}}$ | $C_6^{\text{MB}}$ | $C_7^{\text{MB}}$ | $C_8^{\text{MB}}$ | $C_9^{\text{MB}}$ | $C_{10}^{\text{MB}}$ | Total |
| --- | --- | --- | --- | --- | --- | --- | --- | --- | --- | --- | --- |
| Attractive | 0 | 4 | 0 | 1 | 0 | 5 | 4 | 11 | 11 | 8 | 44 |
| Aversive | 1 | 2 | 0 | 0 | 4 | 12 | 9 | 8 | 8 | 3 | 47 |
| Unknown | 4 | 7 | 8 | 5 | 6 | 5 | 1 | 2 | 3 | 3 | 44 |
| Total | 5 | 13 | 8 | 6 | 10 | 22 | 14 | 21 | 22 | 14 | 135 |

Then the *χ*^2^ value is evaluated based on the table using

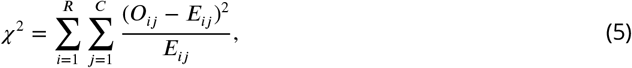

where *R* and *C* are the numbers of rows and columns, and *O*_*ij*_ and *E*_*ij*_ are the observed and expected frequencies of the event in the *i*-th row and *j*-th column, respectively. *E*_*ij*_ is calculated from *O*_*ij*_ as

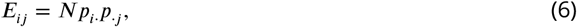

where 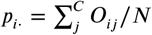 and 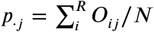 with *N* being the total count. Thus, *E*_*ij*_ is the frequency expected by assuming that the two categorical data are statistically independent. Pearson’s *χ*^2^ test aims to check whether there is a significant difference between *O*_*ij*_ and *E*_*ij*_.

In the *χ*^2^-test, the p-values are estimated using *f*_*k*_(*x*), the *χ*^2^-distribution with the degree of freedom *k* = (*R* − 1)(*C* − 1). If the test returns a *χ*^2^ value that gives rise to a p-value smaller than the defined significance level (*α* = 0.01), the null hypothesis of independence between the two data sets should be rejected. As a result, the distribution of the categorical data is deemed significantly different from a randomly generated distribution, which concludes that the association between two sets of data is statistically significant.

For the above contingency table with *k* = 18, which leads to *χ*^2^ ≈ 66.1 (Equation 5), we get a p-value much smaller than the significance level (*α* = 0.01), 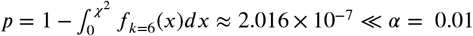.

When Pearson’s *χ*^2^ statistics are available, one can calculate Cramér’s *V* with bias correction, a measure of association between two categorical variables, as follows.

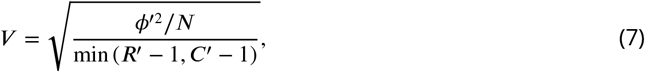

where *ϕ*′^2^ = max (0, *χ*^2^/*N* − (*R* − 1)(*C* − 1)/(*N* − 1)), *R*^*′*^ = *R* − (*R* − 1)^2^/(*N* − 1), and *C*^*′*^ = *C* − (*C* −1)^2^/(*N* − 1). Similar to the Pearson correlation coefficient, the value *V* ranges between 0 and 1 where 0 indicates no correlation and 1 indicates a complete correlation between two categorical variables.

### Mutual information

Mutual information (*I*) is used to verify the significance of association between nominal variables observed in Pearson’s *χ*^2^-test for independence. The *I* measures the information transfer or the similarity between two data. The concept can be extended to clustering outputs to check how two different clustering labels from the same data are similar to each other. Traditionally, the *I* between two jointly discrete variables *A* and *B* is given by

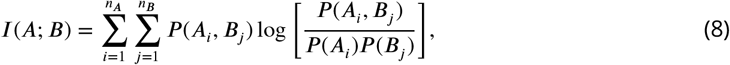

where *n*_*A*_ (or *n*_*B*_) is the number of clusters in *A* (or *B*). Numerically, the between two clustering outputs *A* and *B* is calculated by evaluating 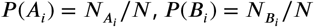, and 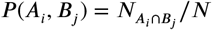 where *N* is the total count and 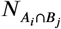 is the number of elements common in both clusters *A*_*i*_ and *B*_*j*_.

The significance was assessed by comparing the observed *I* with the distribution of *I*s from randomly sampled variables. Specifically, the cluster label was randomly sampled 1000 times to generate a distribution of *I* under the assumption of independence. The value of observed *I* is considered significant if the approximated p-value is below 0.01 (*p* < 0.01).

### Analysis of synaptic interfaces

We conducted three different analyses on the synaptic interfaces of PNs with KCs and LHNs.

i. The *‘*homotype-specific*’* connections (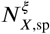 with *ξ* = PN-KC or PN-LHN) are obtained by counting the number of third-order neurons that synapse with a homotype *X* and do not synapse with any other homotypes from the binarized connectivity matrix *C*. The total number of synaptic connections for a homotype *X* is simply the sum of the row of the connectivity matrix *C* 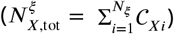.
ii. To generate the *S* matrices, we counted the number of third-order neurons synapsing with a given homotype *X* that also synapses with other homotypes.
iii. The tanglegram study required a hierarchical clustering of uPNs based on their connectivity to third-order neurons. The distances between uPNs in the connectivity matrix *C* are representative of how similar the connectivity patterns to third-order neurons between two uPNs are. We utilized the cosine distance widely used for analyzing the connectivity matrix (***Bates et al., 2020***; ***Li et al., 2020***; ***Schlegel et al., 2021***), which is defined as

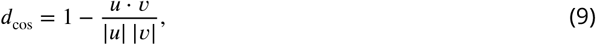

where *u* and *v* are two vectors to be compared. After calculating the distances, we performed hierarchical clustering by Ward’s criterion, which minimizes the variance of merged clusters, to generate the tree structure. The results of hierarchical clustering using the spatial proximity (*d*_*αβ*_) and connectivity (*d*_cos_) are compared using a tanglegram after untangling two trees using the *‘*step1side*’* method (***Galili, 2015***) (Figure 12).

## Data Availability

All data generated during this study and the Python scripts are available in the .zip file included as the supplementary material. They are also available at https://github.com/kirichoi/DrosophilaOlfaction.

## Acknowledgements

We thank Dr. Ji Hyun Bak for helpful discussions. This study was supported by KIAS Individual Grants CG077001 (K.C.), CG076002 (W.K.K.), and CG035003 (C.H.). We thank the Center for Advanced Computation in KIAS for providing the computing resources.

## Declaration of Interests

The authors declare no competing interests.

## Supplementary Information

### Testing the labeled-line hypothesis

We detail the analyses performed on the tanglegram and the respective outputs (Figure 12). First, we applied the dynamic hybrid cut tree method on the dendrogram generated from connectivity and conducted Pearson’s *χ*^2^ test. The results are shown in Table S4. The p-values for the connectivity-based clustering between uPNs and LHNs for glomerular labels, odor types, and odor valence were very small. For the connectivity between uPNs and KCs, we see a moderate to no association for the given categorical variables.

The similarity between two tree structures from spatial proximity-based and connectivity-based clustering at a given synaptic interface is measured in several different ways to provide a comprehensive comparison. First, we quantified the similarity using Baker’s Gamma index (***Baker, 1974***), which is a measure of rank correlation (or ordinal relation) calculated from concordant and discordant pairs given by

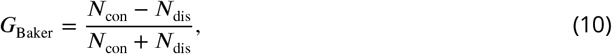

where *N*_con_ is the number of concordant pairs (the ordering of elements in two trees match) and *N*_dis_ is the number of discordant pairs (the ordering of elements in two trees do not match). Baker’s Gamma index ranges from −1 to 1 where 0 represents the ordering of two trees is completely dissimilar and 1 or −1 indicate the ordering of two trees match. We find 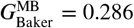 and 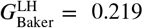 (which we double-checked using both the in-house code and *‘*dendextend*’* package in R). Baker’s Gamma index for LH is very similar to the one obtained by ***Bates et al***. (***2020***) 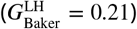, who conducted a similar study using the NBLAST score and connectivity. However, the fact that 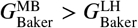 when the tanglegram of MB calyx is seemingly more incoherent (Figure 12A) raises a question of whether Baker’s Gamma index alone is enough to describe the tanglegram.

Apart from the ordinal relations between two sets of leaves, we employed two additional metrics to compare the two trees: (1) entanglement, a measure spanning from 0 to 1 quantifying the number of lines crossing, and (2) cophenetic distance correlation, a measure spanning from 0 to 1 quantifying how similar the two branching structures are. The entanglement between two trees for MB calyx was 0.35 (higher entanglement), while the entanglement for LH was 0.26 (lower entanglement), which agrees with Figure 12. To calculate cophenetic distance correlation, we measured the pairwise cophenetic distances within each tree and calculated the Pearson correlation coefficient. The cophenetic distance between two leaves in the dendrogram is equal to the minimum distance (or height) to the branching point that contains both leaves. The Pearson correlation coefficient between cophenetic distances of the spatial proximity-based and connectivity-based tree structures was *r* = −0.032 (*p* > 0.001) for MB calyx and *r* = 0.236 (*p «* 0.001) for LH, reflecting the less disrupted tree structure in LH compared to MB calyx.

### Analysis based on the hemibrain dataset

The results of this study are mainly based on the FAFB dataset (***Bates et al., 2020***). However, a separate dataset independently reconstructed using a female *Drosophila* is also publicly available (the hemibrain dataset) (***Scheffer et al., 2020***), which allows us to test the generality of our results. One of the major differences between the FAFB dataset and the hemibrain dataset was the number of uPNs that meet our criterion. In the hemibrain dataset, we found 120 uPNs forming 58 glomeruli that innervate all three neuropils according to the annotation provided by the hemibrain dataset. This discrepancy is presumably due to the difference between the neuropil boundary we used and the region defined by the hemibrain dataset. In fact, the total number of uPNs in the two datasets is comparable, with 164 uPNs in the FAFB dataset and 162 uPNs in the hemibrain dataset. The two datasets also had a minor mismatch in the glomerulus label annotations, sometimes affecting the number of uPNs constituting a given homotype.

Figure S8 contains reproductions of several notable figures presented in the main text using the hemibrain dataset. Looking at Figure S8A, we see that 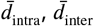, and *λ* are strikingly similar to what is shown in Figure 5A. A minor difference is observed in 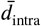 where the hemibrain dataset seems to have a slightly tighter bundling structure. In Figure S8B, we see small differences in 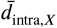 and 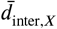 compare to the output of the FAFB dataset due to the discrepancy discussed above, which is expressed as changes to the individual *λ*_*X*_ values, the ordering of homotypes, and the list of single uPN homotypes. However, the overall output is quite similar, which translates to a similar Pearson correlation coefficient (*r* = 0.677, *p* < 0.0001) between *λ*_*X*_ s in MB calyx and LH as shown in Figure S8C, a result that is very close to Figure 7C. Hierarchical clustering following the same methodology resulted in 14 clusters for uPN innervation in MB calyx and 13 clusters in LH. Here, the overall results observed from the FAFB dataset are well-translated and a comparable clustering output is obtained (Figure S8D), where many homotypes clustered together in the FAFB dataset are also clustered in the hemibrain dataset. The spatial segregation of pheromone-encoding uPNs, hygro/thermo-sensing uPNs, and the mixture of uPNs encoding food-related odors are still present. For example, the majority of pheromone-encoding uPNs are sorted into the cluster 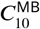 from the hemibrain dataset, which is comparable to the cluster 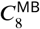 from the FAFB dataset (Figure 8). Hygro/thermo-sensing uPNs are generally segregated from other odor-encoding uPNs in the hemibrain dataset. In LH, many uPNs encoding aversive responses are clustered together (e.g. clusters 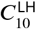 and 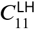 in Figure S8D) and many food-related homotypes are clustered together (e.g. clusters 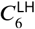 and 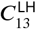 in Figure S8D). Also, various statistical tests (e.g., Pearson’s *χ*^2^-test) reached the same conclusion using the hemibrain dataset (data not shown). Overall, we report that our analysis using the hemibrain dataset faithfully reproduced the major outcomes of our results based on the FAFB dataset.

### Monte Carlo approach to independence test

In this section, we describe an alternative method to the independence test inspired by the Monte Carlo significance test (***Hope, 1968***) to further support our Pearson’s *χ*^2^-test of independence. The procedure is as follows: 1) For a given contingency table, randomize the observation such that the marginal sum of each row remains the same as the observed contingency table. That is, for each row, randomize the vector with integers while the sum of the vector stays the same as the observed contingency table. This procedure randomly shuffles the distribution of the clusters while keeping the distribution of a particular categorical variable intact. 2) Calculate the *χ*^2^ value from the randomized contingency table. 3) Repeat steps 1 and 2 for 1000 times to generate a distribution of the *χ*^2^ values. 4) Obtain the mean and the standard deviation of *χ*^2^ values. The distribution of *χ*^2^ values is approximately normal. 5) If the *χ*^2^ value from the observed contingency table is more than 4σ different from the distribution, we consider the observed *χ*^2^ value statistically significant and reject the null hypothesis. Whenever we ran a Pearson’s *χ*^2^-test, we performed the above procedure alongside. The output of this procedure supported whichever conclusion we drew from Pearson’s *χ*^2^-test.

### Identifying the agreement between two categorical data via mutual information

We verified our Pearson’s *χ*^2^-test of independence of two categorical variables by calculating the mutual information *I* (see Methods). The mutual information between glomerular labels and *d*_*αβ*_ -based clustering output in MB calyx was equal to *I* (glo; *C*^MB^) = 1.892, which is significantly (more than 4 σ) different from the mean of randomly sampled distribution (*I* (glo; *C*^MB^)_rand_ = 1.386 ± 0.035). This result is consistent with our *χ*^2^-test, as the mutual information of the observed variables is significantly larger than the mutual information under the assumption of random sampling, suggestive of a statistically significant association between glomerular labels and MB calyx cluster labels. In LH, the mutual information between glomerular labels and *d*_*αβ*_ -based clustering output was *I* (glo; *C*^LH^) = 2.128 which deviated 4*σ* or more from the mean of the randomly sampled *I* distribution (*I* (glo; *C*^LH^)_rand_ = 1.466 ± 0.035).

The same method is applied to confirm that a statistically significant association exists between odor type and the clustering outputs, with *I* (odor; *C*^MB^) = 0.819 and (odor; *C*^LH^) = 0.963, all of which differ by more than 4 σ from the means of the randomly sampled *I* distributions (*I* (odor; *C*^MB^)_rand_ = 0.337 ± 0.044, *I* (odor; *C*^LH^)_rand_ = 0.372 ± 0.043). For odor valence, we obtain *I* (val; *C*^MB^) = 0.277 and *I* (val; *C*^LH^) = 0.326, where both *I* (val; *C*^MB^) and *I* (val; *C*^LH^) differ significantly from the means of the randomly sampled *I* distributions (*I* (val; *C*^MB^)_rand_ = 0.073 ± 0.026, *I* (val; *C*^LH^)_rand_ = 0.081 ± 0.026). Overall, the conclusion drawn from the association study based on mutual information is identical to Pearson’s *χ*^2^-test.

**Table S1.** Pearson’s *χ*^2^ tests of independence of variables. *C*^*Z*^ indicates cluster labels from *d*_*αβ*_ -based clustering in *Z* neuropil. Cramér’s V values are displayed on each cell and the corresponding p-values are shown in parentheses. Bold entries are used to specify statistically significant results.

| | $C^{\text{LH}}$ | Glomerular Labels | Odor Type | Odor Valence |
| --- | --- | --- | --- | --- |
| $C^{\text{MB}}$ | **0.502**<br>(1.149E-36) | **0.610**<br>(1.255E-27) | **0.401**<br>(3.303E-21) | **0.425**<br>(2.016E-07) |
| $C^{\text{LH}}$ | | **0.671**<br>(2.266E-40) | **0.416**<br>(1.980E-22) | **0.455**<br>(2.586E-08) |

**Table S2.** Mutual information (observed mutual information (top), randomly sampled mutual information (bottom) in each cell) from the association study. *C*^*Z*^ is cluster labels from *d*_*αβ*_-based clustering at *Z* neuropil. Bold entries are used for observed *I* that are more than 4σ different from the randomly sampled *I*.

| | $C^{\text{LH}}$ | Glomerular Labels | Odor Type | Odor Valence |
| --- | --- | --- | --- | --- |
| $C^{\text{MB}}$ | **1.076**<br>$0.397 \pm 0.045$ | **1.892**<br>$1.386 \pm 0.035$ | **0.819**<br>$0.337 \pm 0.044$ | **0.277**<br>$0.073 \pm 0.026$ |
| $C^{\text{LH}}$ | | **2.128**<br>$1.466 \pm 0.035$ | **0.963**<br>$0.372 \pm 0.043$ | **0.326**<br>$0.081 \pm 0.026$ |

**Table S3.** Statistics of homotypes composed of a single uPN (or multiple uPNs) and the corresponding putative valence.

|  | Aversive | Attractive | Unknown | Total |
| --- | --- | --- | --- | --- |
| Single uPN Homotypes Count | 7 | 4 | 2 | 13 |
| Multiple uPN Homotypes Count | 18 | 13 | 13 | 44 |
| Total | 25 | 17 | 15 | 57 |

**Table S4.** Pearson’s *χ*^2^ tests of independence of variables on the connectivity-based clustering results. Cramér’s V values are displayed on each cell and the corresponding p-values are shown in parentheses. Bold entries are used to specify statistically significant results.

|  | Glomerular Labels | Odor Type | Odor Valence |
| --- | --- | --- | --- |
| $C^{\text{PN-KC}}$ | **0.433**<br>(2.472E-08) | **0.316**<br>(9.978E-09) | 0.271<br>(0.012) |
| $C^{\text{PN-LHN}}$ | **0.765**<br>(1.410E-67) | **0.630**<br>(1.519E-48) | **0.604**<br>(4.055E-12) |

**Figure S1.**
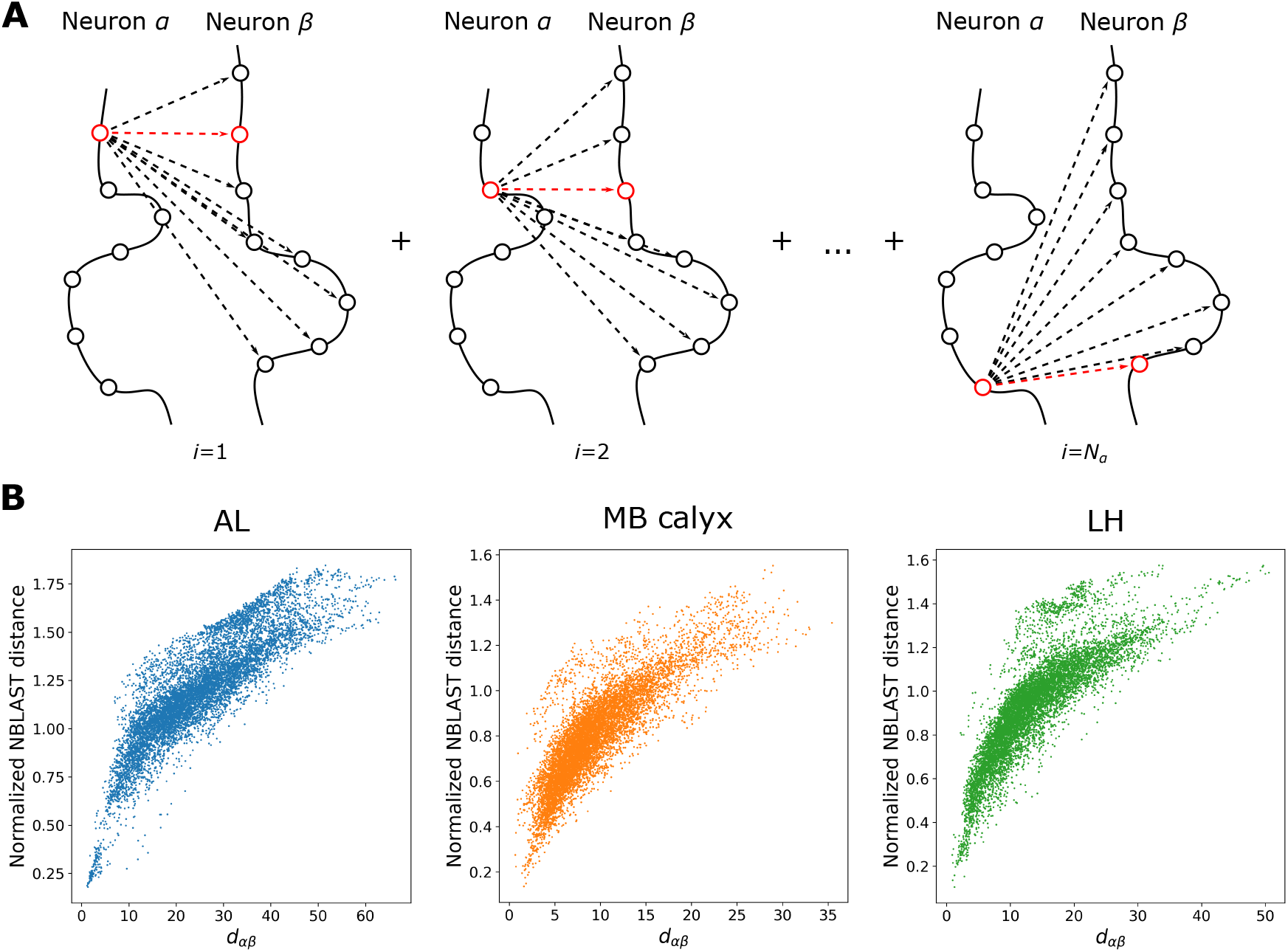
(A) A schematic showing how the *‘*distance*’* between two neurons *α* and *β* (*d*_*αβ*_) is calculated. (B) The comparisons between *d*_*αβ*_ and the normalized NBLAST distance (which is equal to 1 − NBLAST score) for uPN innervation to AL, MB calyx, and LH. While the two distances are correlated, significant dispersion is also present. Unlike NBLAST score, *d*_*αβ*_ measures the spatial proximity between two neurons *α* and *β* only, but not their morphological similarity.

**Figure S2.**
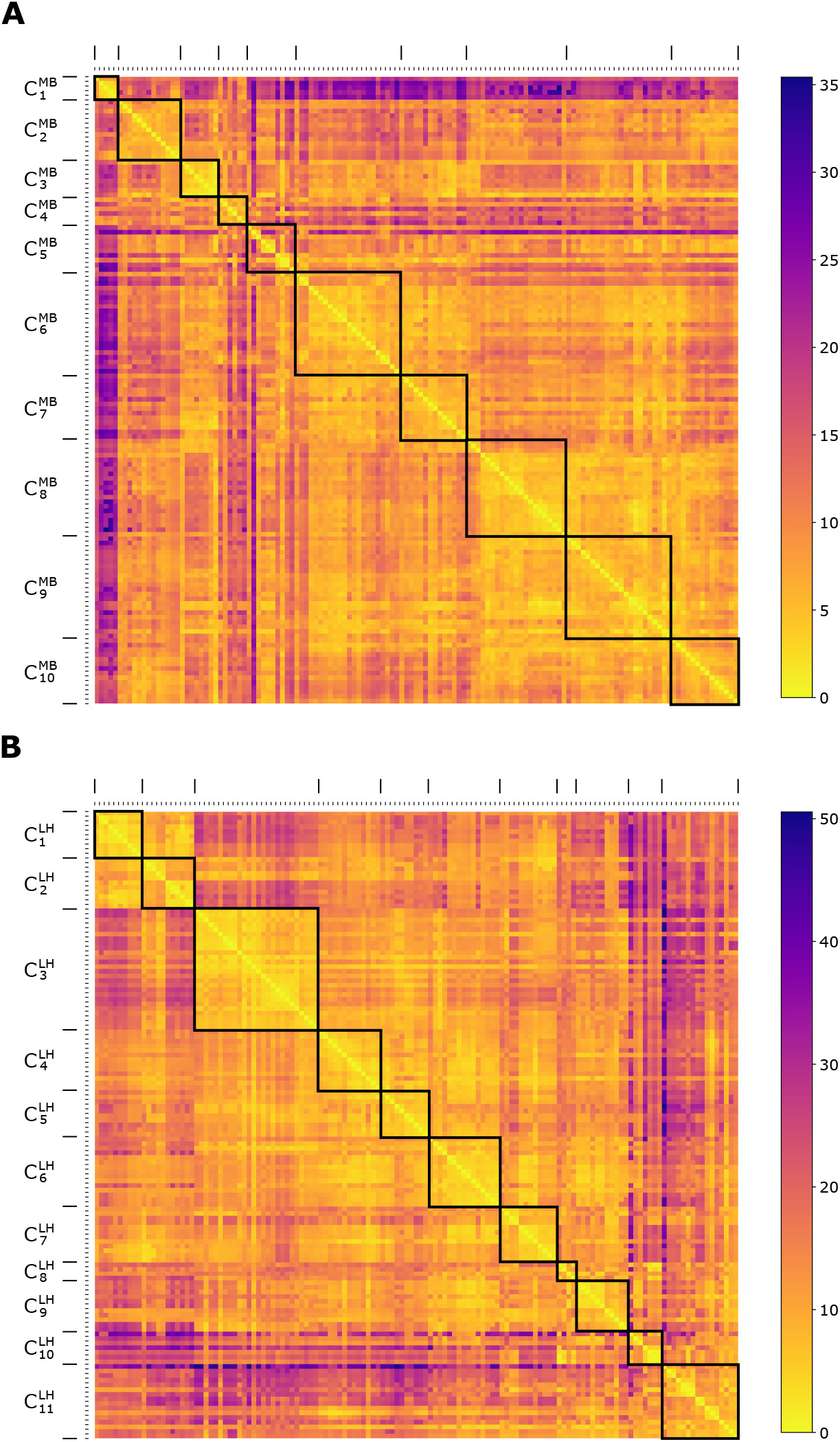
Two 135×135 matrices representing the inter-neuronal distances in units of *µ*m between individual uPN innervation in (A) MB calyx and (B) LH. The diagonal block represents each cluster (see Figures 3, 4, and S3 for detailed information on the clustering labels). The indices of uPN in Figure S2 are reorganized based on the results from the clustering analysis.

**Figure S3.**
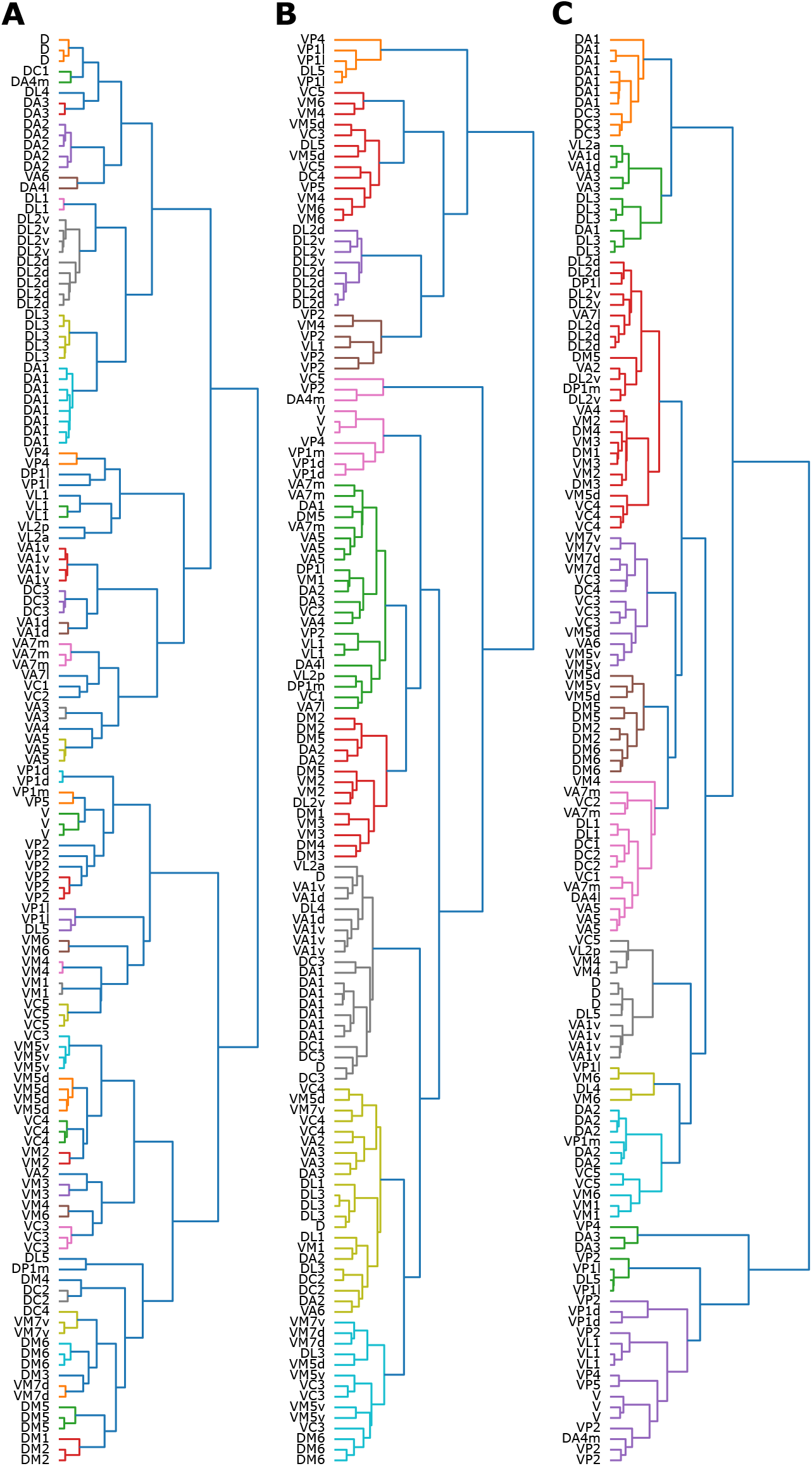
The dendrograms of *d*_*αβ*_ -based clustering on uPNs innervation in (A) AL, (B) MB calyx, and (C) LH. In (B) and (C), the different colored leaves correspond to a cluster generated from the tree-cutting method. A leaf represents an individual uPN and the label depicts the corresponding homotype for each uPN.

**Figure S4.**
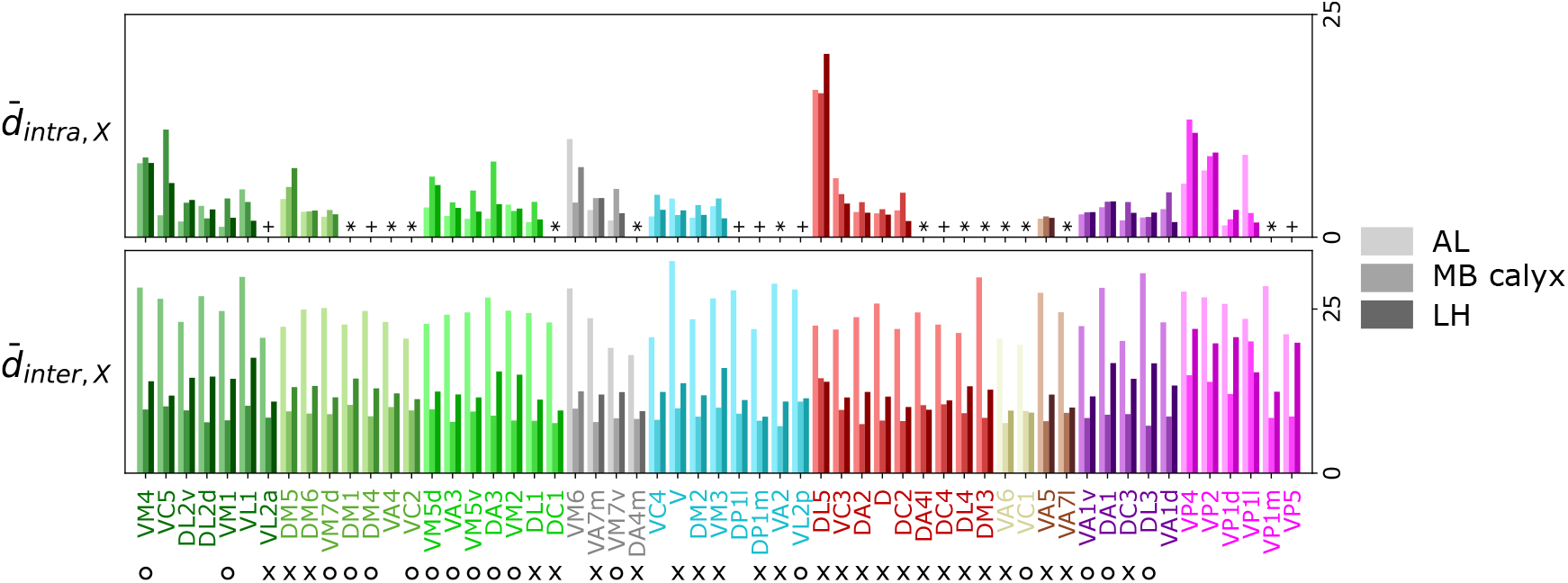
Comparison of the intra- (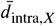, degree of bundling) and inter-PN (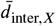, degree of packing) distances of *X*-th homotype in AL, MB calyx, and LH (from lighter to darker colors). The homotype label is color-coded based on the odor types obtained from the literature (Dark green: decaying fruit, lime: yeasty, green: fruity, gray: unknown, cyan: alcoholic fermentation, red: general bad/unclassified aversive, beige: plant matter, brown: animal matter, purple: pheromones, pink: hygro/thermo). Homotypes are ordered based on both the odor type and the values of *λ*_*X*_ in LH. Asterisks (*) mark homotypes composed of a single uPN while plus (+) mark homotypes composed of a single uPN under our selection criterion but are actually a multi-uPN homotype, whose intra-homotype uPN distance is not available. O and × denote attractive and aversive odors, respectively.

**Figure S5.**
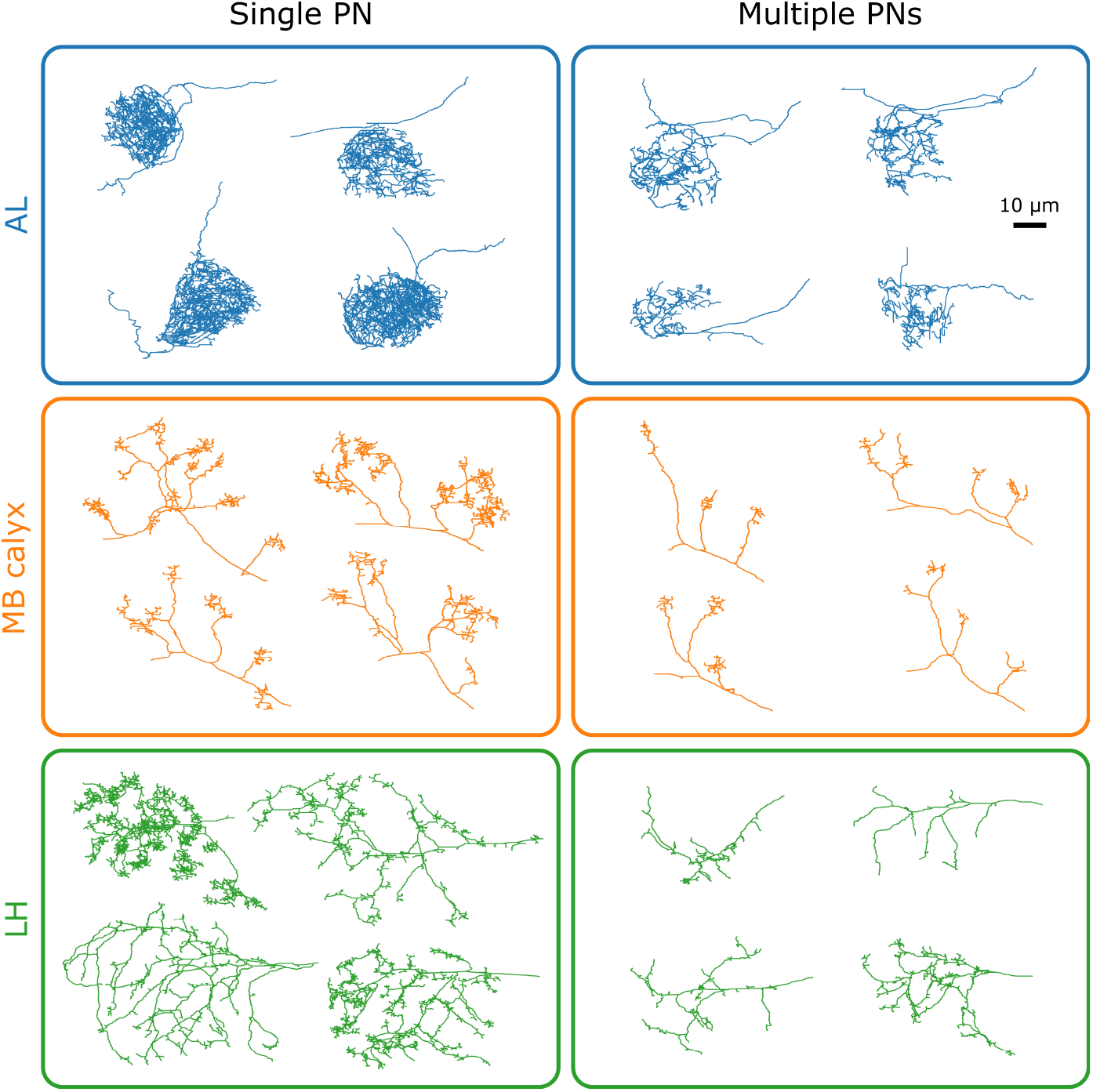
Selected morphologies of uPN innervation at each neuropil for single uPN homotypes and multiple uPN homotypes.

**Figure S6.**
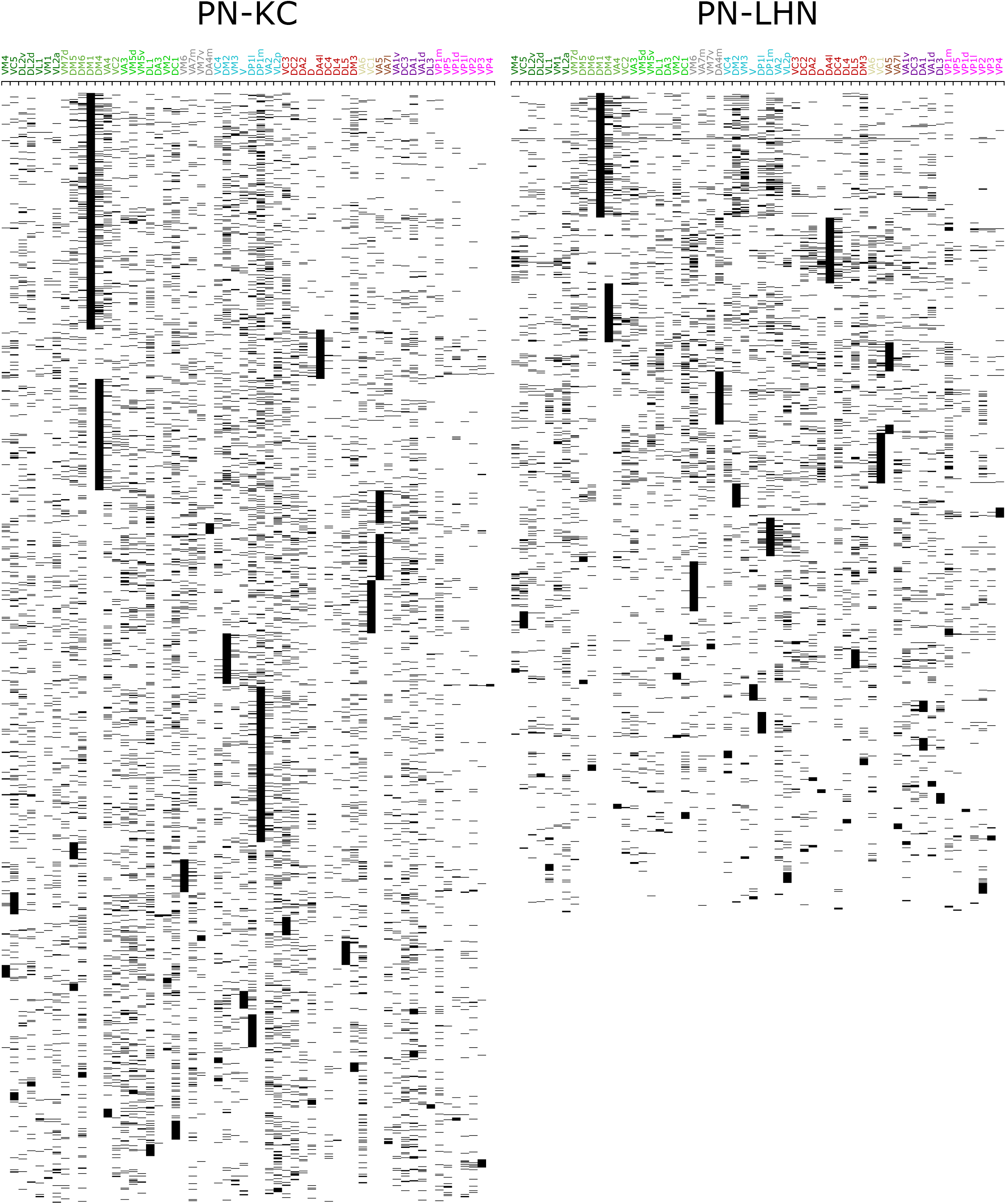
The 58 × 1754 and 58 × 1285 synaptic connectivity matrices for the synaptic interface between PNs and KCs (*C*^PN−KC^) (left) and between PNs and LHNs (*C*^PN−LHN^) (right). Homotypic uPNs are merged together.

**Figure S7.**
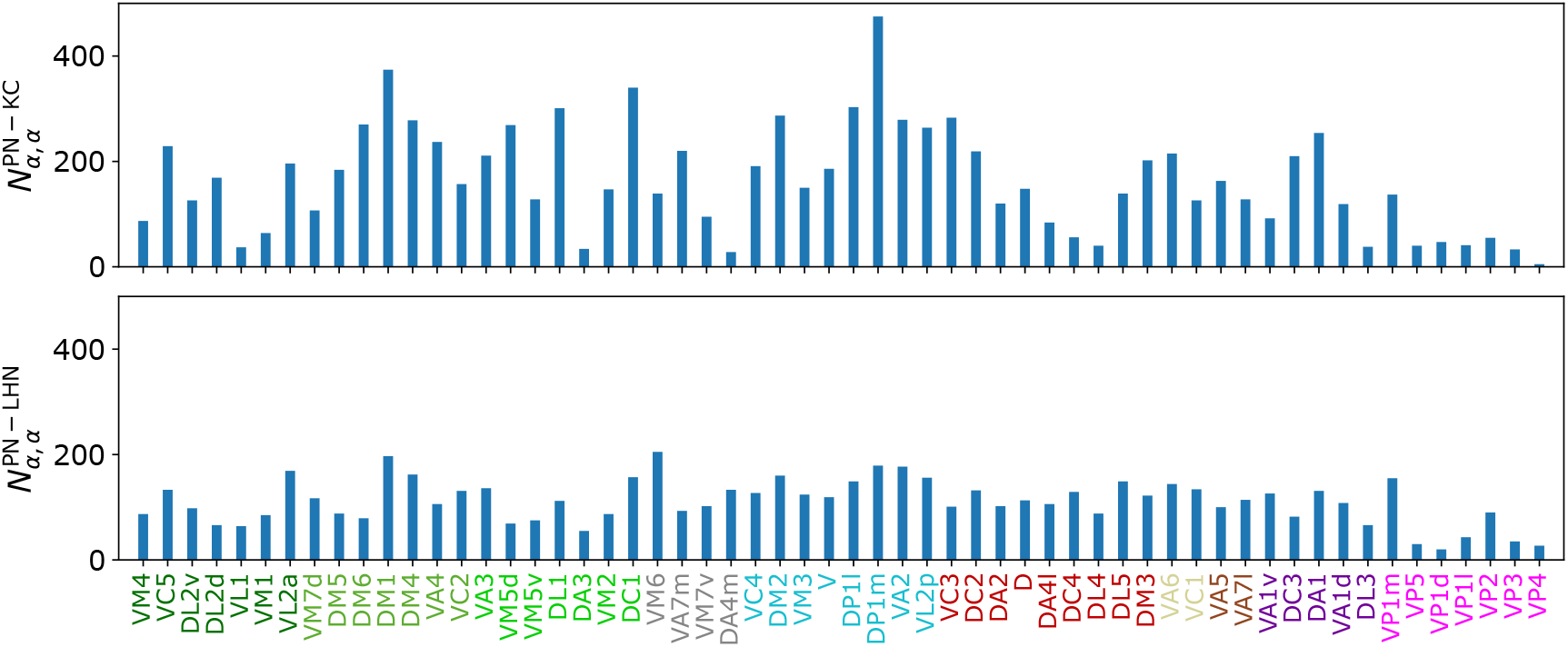
The total number of third-order neurons connected to each homotype, which corresponds to the diagonal element of common synapse matrix (*S*^*ξ*^) in MB calyx (top) and LH (bottom).

**Figure S8.**
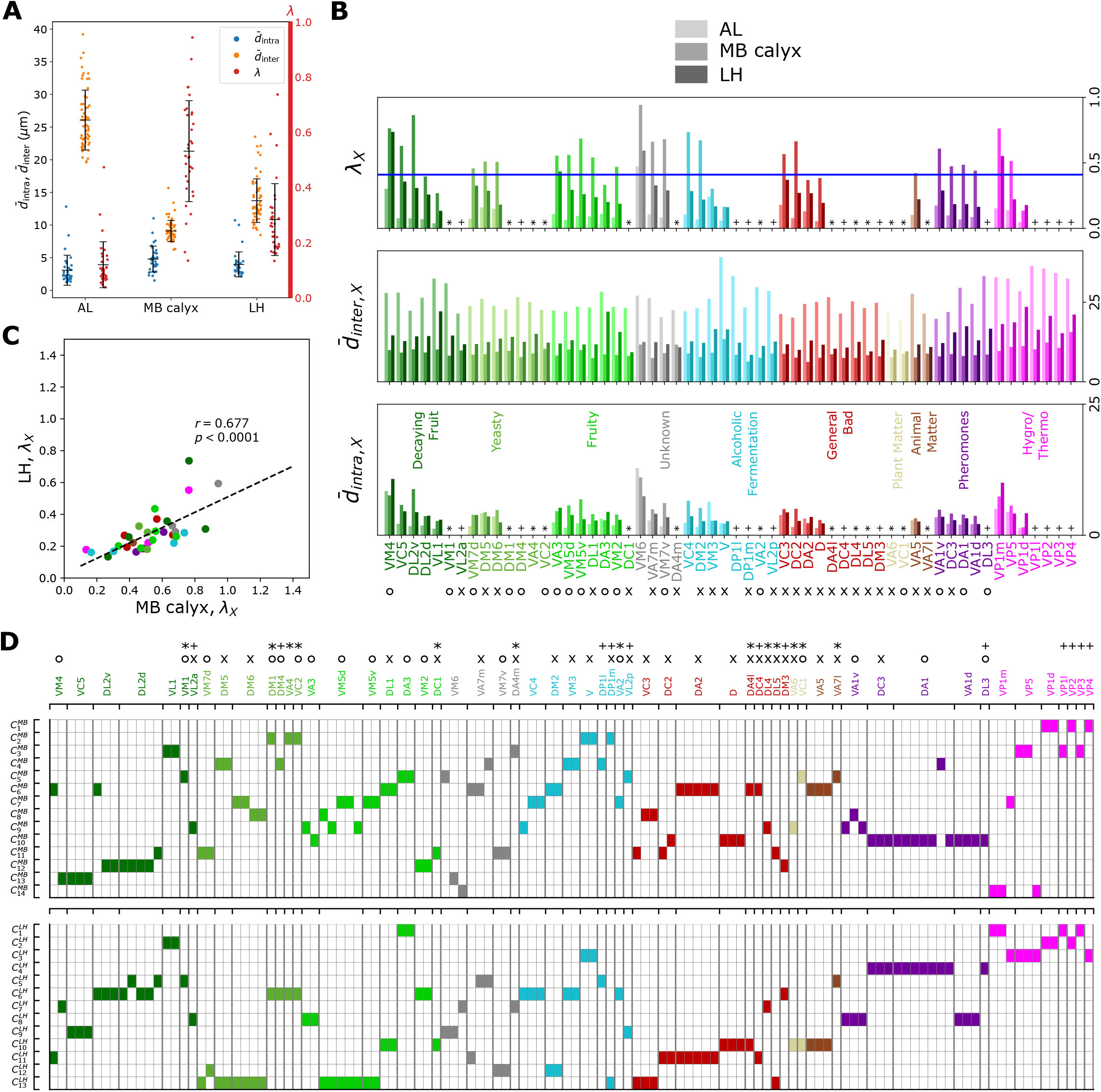
Analyses of hemibrain dataset. (A) A graph depicting 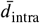 (blue, degree of bundling), 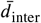 (orange, degree of packing), and *λ* (red, degree of overlapping), which can be compared with Figure 5A. (B) The degree of overlapping (*λ*_*X*_) for *X*-th homotype in AL, MB calyx, and LH (from lighter to darker colors), which can be compared with Figures 6A and S4. (C) Scatter plot depicting the relationships between *λ*_*X*_s in LH versus *λ*_*X*_s in MB calyx, which can be compared with Figure 7C. (D) A diagram summarizing how the clusters of uPN innervation in MB calyx (14 clusters) and LH (13 clusters) are associated with the odor types, which can be compared with Figure 8.

**Figure S9.**
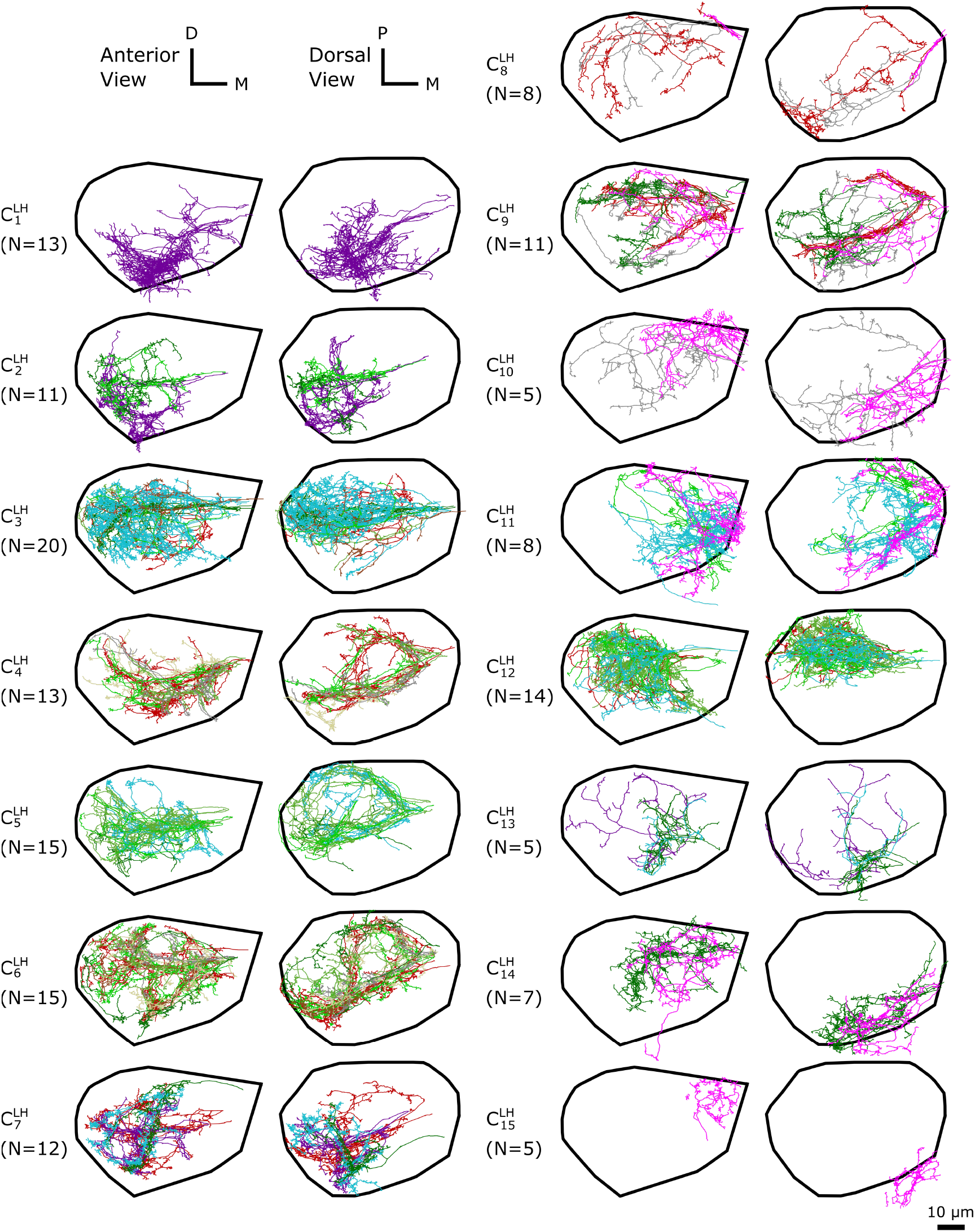
The *d*_*αβ*_-based clustering on the entire uPN innervation in LH resulting in 15 clusters. The individual uPNs are color-coded based on the encoded odor types (Dark green: decaying fruit, lime: yeasty, green: fruity, gray: unknown/mixed, cyan: alcoholic fermentation, red: general bad/unclassified aversive, beige: plant matter, brown: animal matter, purple: pheromones, pink: hygro/thermo). The first and second columns illustrate the anterior and the dorsal view, respectively (D: dorsal, M: medial, P: posterior). The black line denotes the approximate boundary of LH.

**Figure S10.**
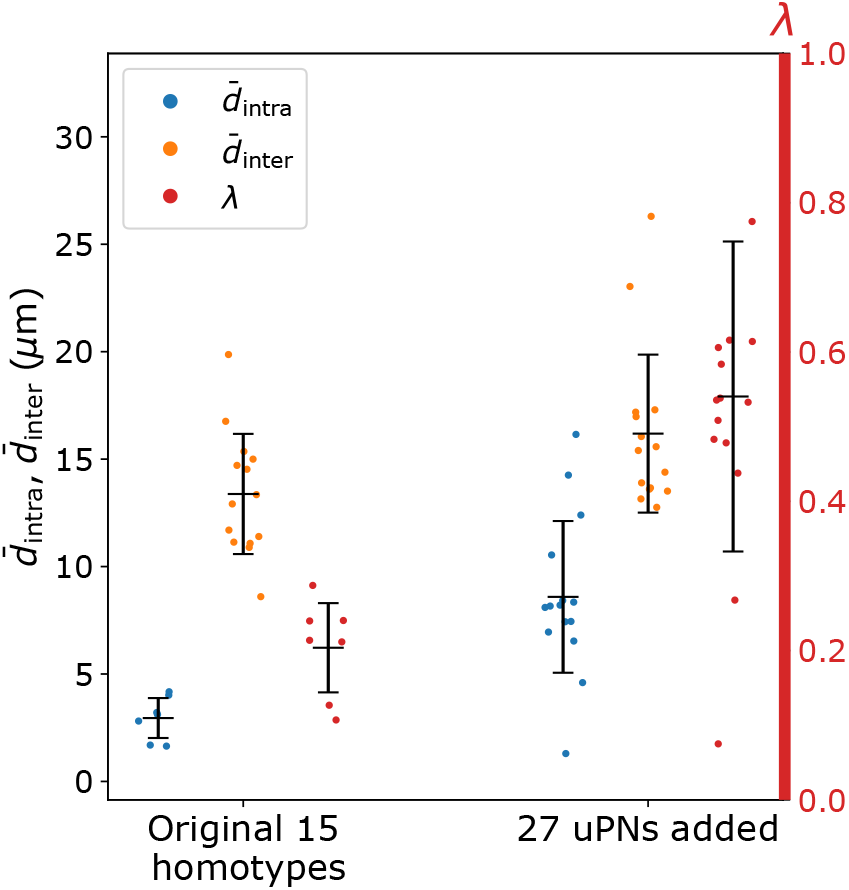
A graph depicting 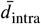 (blue, degree of bundling), 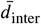 (orange, degree of packing), and the ratio between the two distances *λ* (red, degree of overlapping) of 15 homotypes without (left) and with (right) 27 uPNs added, which are mostly GABAergic and follow mlALT.

**Figure S11.**
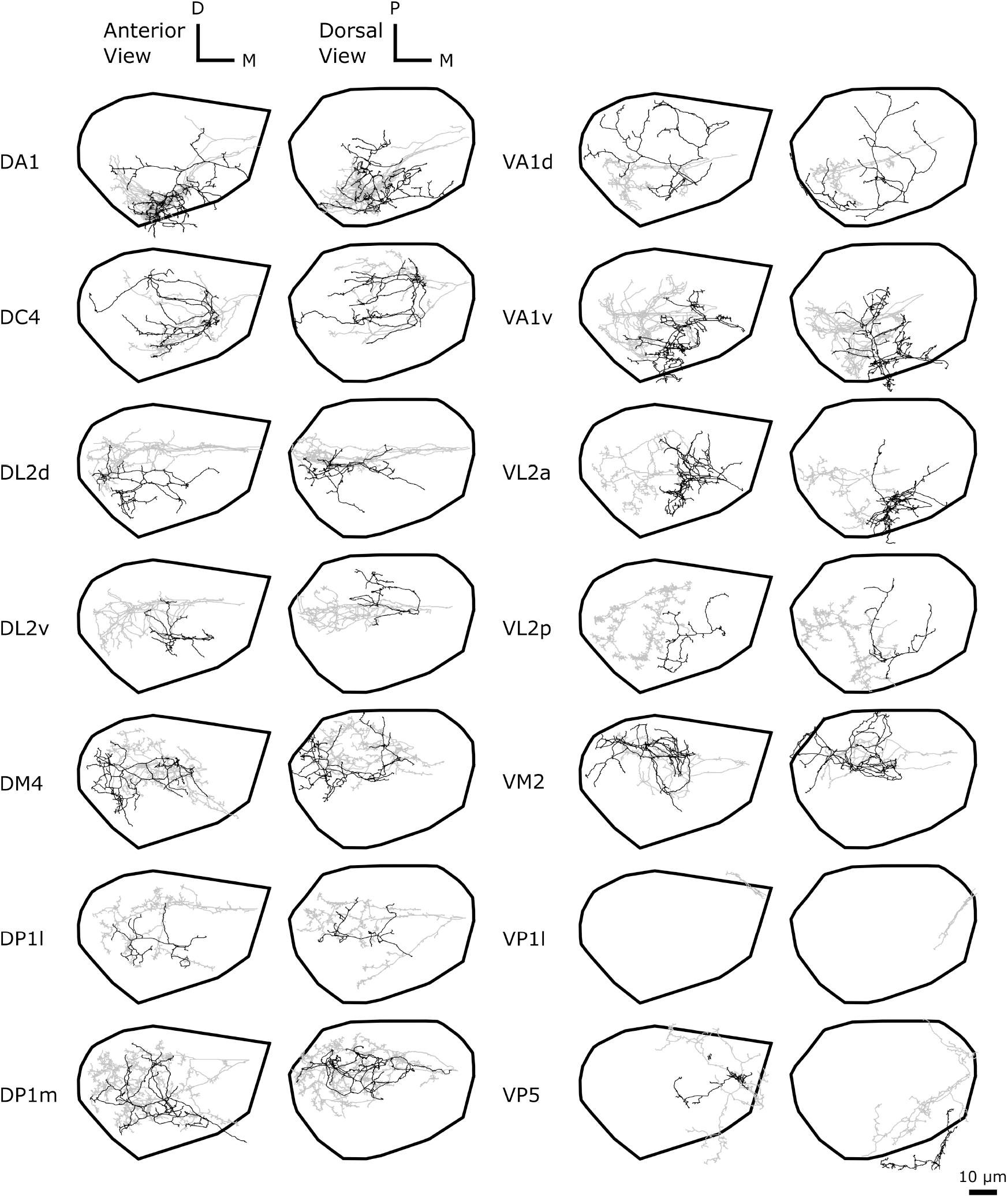
Comparison of innervation pattern in LH between the uPNs innervating all three neuropils (gray, most of which follow mALT) and the local uPNs (black, most of which follow mlALT). Shown are 14 homotypes consisting of uPNs whose innervation is localized to LH.

**Figure S12.**
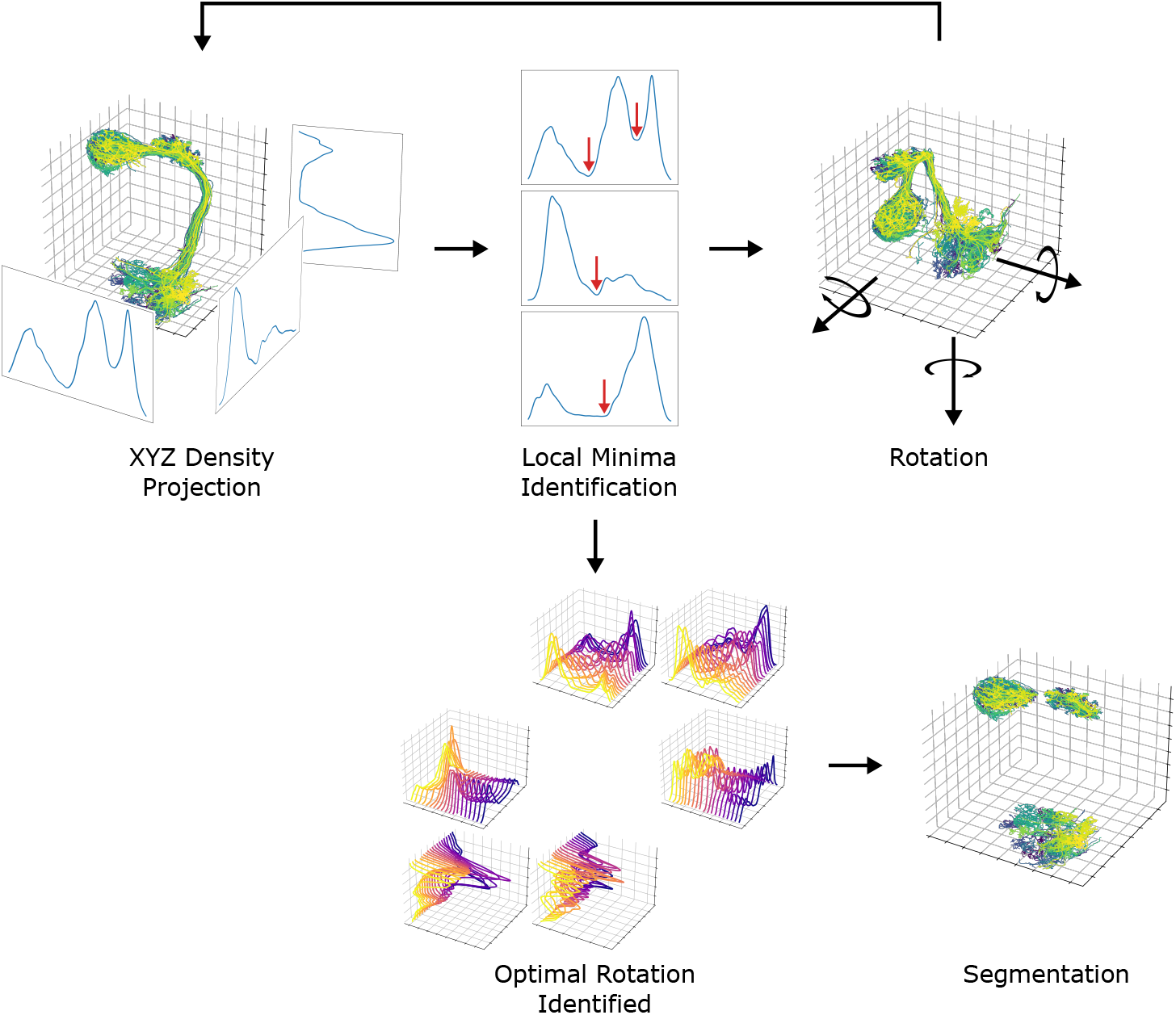
A diagram depicting the neuropil segmentation process. The data points from skeletal reconstruction are projected to each axis to generate distributions from which local minima are obtained. The process is repeated while rotating the uPNs along each axis. A collection of histograms and corresponding local minima are surveyed to generate a set of optimal rotations and boundaries for individual neuropil. The resulting parameters are combined to form a collection of conditions to segment each neuropil.

## Notes

### Competing Interest Statement

The authors have declared no competing interest.

### Summary of Updates

Figures 8-12 and the corresponding text were newly added.

